# Neuronal Modeling of Alternating Hemiplegia of Childhood Reveals Transcriptional Compensation and Replicates a Trigger-Induced Phenotype

**DOI:** 10.1101/2020.04.08.031732

**Authors:** John P. Snow, Grant Westlake, Lindsay K. Klofas, Soyoun Jeon, Laura C. Armstrong, Kathryn J. Swoboda, Alfred L. George, Kevin C. Ess

## Abstract

Alternating hemiplegia of childhood (AHC) is a rare neurodevelopmental disease caused by heterozygous de novo missense mutations in the ATP1A3 gene that encodes the neuronal specific α3 subunit of the Na,K-ATPase (NKA) pump. Mechanisms underlying patient episodes including environmental triggers remain poorly understood, and there are no empirically proven treatments for AHC. In this study, we generated patient-specific induced pluripotent stem cells (iPSCs) and isogenic controls for the E815K ATP1A3 mutation that causes the most phenotypically severe form of AHC. Using an in vitro iPSC-derived cortical neuron disease model, we found elevated levels of ATP1A3 mRNA in AHC lines compared to controls, without significant perturbations in protein expression. Microelectrode array analyses demonstrated that in cortical neuronal cultures, ATP1A3+/E815K iPSC-derived neurons displayed a non-significant trend toward less overall activity than neurons differentiated from isogenic mutation-corrected and unrelated control cell lines. However, induction of cellular stress by elevated temperature revealed a hyperactivity phenotype following heat stress in ATP1A3+/E815K lines compared to control lines. Treatment with flunarizine, a drug commonly used to prevent AHC episodes, did not impact this stress-triggered phenotype. These findings support the use of iPSC-derived neuronal cultures for studying complex neurodevelopmental conditions such as AHC and provide a potential route toward future therapeutic screening and mechanistic discovery in a human disease model.

## 1. Introduction

Alternating hemiplegia of childhood (AHC) is a rare and devastating neurodevelopmental disorder caused by heterozygous missense mutations in *ATP1A3*, the gene encoding the α3 subunit of the Na,K-ATPase (NKA) ion transporter (Heinzen et al., 2012). The NKA complex is responsible for exchanging three cytoplasmic sodium ions for two extracellular potassium ions, thus establishing electrochemical gradients essential for crucial cellular functions and membrane potential (Benarroch, 2011; Clausen et al., 2017; Dobretsov and Stimers, 2005).

Patients with AHC present early in life with a constellation of symptoms including abnormal eye movements, seizures, dystonia, and characteristic spells of alternating hemiplegia or full quadriplegia (Bourgeois et al., 1993; Heinzen et al., 2014). Episodes in AHC are often triggered by external stimulation including temperature changes, physical exertion, or exposure to certain foods, among other triggers (Kansagra et al., 2013; Sweney et al., 2009). Over time, AHC patients manifest developmental delay and acquire intellectual disabilities.

AHC is the best studied to date of an expanding spectrum of neurological conditions associated with *ATP1A3* mutations (Sweney et al., 2015), including RDP (rapid-onset dystonia parkinsonism), CAPOS (cerebellar ataxia, areflexia, pes cavus, optic nerve atrophy, and sensorineural hearing loss), RECA (relapsing encephalopathy with cerebellar ataxia), EIEE (early infantile epileptic encephalopathy), and FIPWE (fever-induced paroxysmal weakness and encephalopathy) (de Carvalho Aguiar et al., 2004; Demos et al., 2014; Heinzen et al., 2014; Paciorkowski et al., 2015; Rosewich et al., 2014). While these syndromes share symptomatic profiles including triggered episodes, a strong genotype-to-phenotype correlation exists within this spectrum. In fact, 60% of patients worldwide diagnosed clinically as AHC have one of three unique *ATP1A3* missense mutations, leading to amino acid changes D801N, E815K, or G947R (Heinzen et al., 2012; Viollet et al., 2015). Only symptomatic treatment options exist for patients diagnosed with AHC, including benzodiazepines and other antiepileptic drugs. A non-specific sodium and calcium channel blocker, flunarizine, has been anecdotally used to treat many patients for the past several decades (Casaer, 1987). Flunarizine has been reported in several recent studies to reduce the severity, duration, or frequency of hemiplegic spells in some patients with AHC, although not all individuals respond to treatment (Delorme et al., 2017; Kansagra et al., 2013; Pisciotta et al., 2017). Conclusions from these studies are limited by small numbers of patients and lack of randomized control groups.

While the NKA-α1 subunit is ubiquitously expressed across cell types, expression of the NKA-α3 subunit is generally restricted to neurons (Clausen et al., 2017; Richards et al., 2007). A critical component of neuronal functionality, the α3 subunit has lower affinity for sodium ions allowing the rapid expulsion of positive ions following neuronal activity and depolarization, particularly in dendrites (Azarias et al., 2013; Blom et al., 2011; Kim et al., 2007). Common AHC-causing mutations in the α3 subunit result in impaired ion transporting ability without impacting protein trafficking to the plasma membrane (Heinzen et al., 2014; Koenderink et al., 2003; Li et al., 2015). A recent study demonstrated lower pump current and depolarized resting membrane potential in AHC-mutant iPSC-derived excitatory neurons (Simmons et al., 2018), and *Drosophila* and mouse models of *Atp1a3* mutations associated with human disease often replicate trigger-induced symptoms or exacerbation (DeAndrade et al., 2011; Helseth et al., 2018; Holm and Lykke-Hartmann, 2016; Isaksen et al., 2017; Palladino et al., 2003; Sugimoto et al., 2014).

In this study, we created a patient-specific model of AHC to determine if a common and phenotypically severe mutation, E815K, results in transcriptional or protein level NKA subunit compensation for impaired pump function in iPSC-derived neurons. Using both unrelated and isogenic corrected *ATP1A3* controls, we measured transcript levels of multiple NKA alpha and beta subunits. We found elevation of *ATP1A3* mRNA during neuronal differentiation of iPSCs, while α3 protein expression showed a nonsignificant trend towards higher levels. We then studied heat-stress within this model employing microelectrode array (MEA) analyses to determine if a trigger-induced phenotype was replicated in this system. We found that rapid temperature increase resulted in diminished neuronal firing in both control and AHC patient neurons. However, neurons derived from AHC patient lines revealed an exaggerated rebound of activity and ultimately became hyperactive compared to controls. Treatment with flunarizine, the non-FDA approved drug used anecdotally to prevent AHC patient episodes, had minimal impact on neuronal firing and temperature-triggered rebound hyperactivity. Our findings importantly suggest that elevated expression of *ATP1A3* mRNA may be both a feature of AHC pathogenesis and a possible biomarker for future *in vitro* studies. Additionally, modeling complex symptoms such as trigger-induced episodes is possible in iPSC-derived neuronal models, providing a potential route for screening of future treatments including both small molecules and gene replacement or genome editing therapies.

## 2. Materials & methods

### 2.1. Study subjects

Cell lines from a subject diagnosed with AHC were generated for this study. AHC patient 91759 is female and diagnosed with AHC at 2 years of age. She first displayed symptoms at 4 weeks of age including tonic arm extension and abnormal ocular movements. Spells of alternating hemiplegia lasting 1-2 days began at 5 months of age, with quadriplegic episodes starting at 10 months. Genetic testing subsequently revealed a heterozygous c.2443G>A (p.E815K) missense mutation in the *ATP1A3* gene. The patient had subsequent disease progression including developmental delay, intellectual disability, and behavioral issues. Two iPSC clones generated from this patient line were used in this study (AHC-20, AHC-24). Isogenic wildtype controls (IC1, IC2) were then created from CRISPR/Cas9 editing of patient clone AHC-24 as described below. Unaffected female (UC1 [CC3], 18 year old) and male (UC2 [CX3], 25 year old) unrelated controls were also used in this study and previously reported (Armstrong et al., 2017; Kumar et al., 2014).

### 2.2 Generation and validation of iPSCs

Fibroblasts were isolated by 3 mm punch skin biopsy from unrelated control volunteers or the described AHC patient following informed consent of volunteer or parent/guardian (study approved by the Vanderbilt Institutional Review Board). Isolated fibroblasts were cultured in DMEM (Gibco) with 10% FBS, 1% non-essential amino acids, and 1% penicillin-streptomycin. To reprogram fibroblasts to pluripotency, dividing cells were electroporated using the Neon System (Invitrogen) with plasmids expressing *OCT4, SOX2, KLF4, L-MYC, and LIN28* (Addgene plasmids #27077, #27078, and #27080) using previously published protocols (Armstrong et al., 2017; Okita et al., 2011). Two days following transfection, culture media was changed to TeSR-E7 (StemCell Technologies). Defined iPSC-like colonies were manually isolated 3-4 weeks post-transfection and expanded on Matrigel (Corning) in mTeSR1 (StemCell Technologies). Validation was performed through: pluripotency marker validation (Nanog, Oct4, SSEA3, SSEA4, and TRA-1-60); embryoid body differentiation and immunostaining with trilineage tissue markers defining ectoderm (Sox1, β-III-Tubulin), mesoderm (smooth muscle actin), and endoderm (GATA4, Sox17); and karyotype analysis of iPSC clones (Genetic Associates Inc, Nashville, TN). See supplementary reagents Table for additional information on antibodies, dilutions, and applications. Validated iPSCs were cultured on Matrigel-coated plates, fed daily with mTeSR1, and passed weekly at a ratio of 1:10-50 dependent on colony confluency using ReLeSR (StemCell Technologies).

### 2.3 Creation of isogenic wildtype lines and genotyping

Genome editing was performed following previously published methods (Ran et al., 2013). CRISPR guide RNAs were designed to target the region containing the AHC-causing *ATP1A3* c.2443G>A mutation in exon 18 using an online tool (ChopChop). Single-stranded DNA oligos were designed to flank the targeted cut site by 60-80 bps on each side and contained both the intended wildtype correction along with synonymous PAM-site mutations to limit repetitive cutting events. sgRNA oligos (IDT) were phosphorylated, annealed, and ligated into plasmid DNA containing the Cas9 enzyme and a puromycin resistance cassette (PX459, Addgene) followed by exonuclease treatment. Ligated plasmids were transformed into DH5-alpha competent *E. coli*. Plasmid DNA was sequence verified using a primer for the U6 promoter. iPSCs were dispersed to single cells with Accutase (StemCell) and suspended to a concentration of 10^6^ cells per 100 μL. Plasmid constructs were delivered to suspended solutions of iPSCs containing the AHC patient genotype *ATP1A3^E815K/+^* genotype by electroporation (Neon System, settings: 1200 V, 20 ms, 2 pulses) and plated at low density in mTeSR1 with 10 μM ROCK-inhibitor Y-27632 (StemCell). Thirty-six hours after plasmid delivery, selection for plasmid expression in iPSCs was performed by a 48-hour treatment of 0.5 μg/mL puromycin. mTeSR1 media was subsequently changed every other day for the following two weeks. Surviving colonies were picked and expanded for genotyping and continued culture. PCR amplification for genotyping was performed using primers described in the supplemental methods. Restriction digest with BsiWI-HF (NEB, R3553) followed by confirmatory sequencing of products (GenHunter) identified corrected clones. Confirmed isogenic wildtype iPSC clones were validated for pluripotency with methods described above. Deep sequencing of line IC2 (isogenic corrected) was also done using the VANTAGE Core at Vanderbilt to validate the homozygous genotype. The expected synonymous PAM-site change (115 variant reads, 0 reference reads) was found in addition to the absence of original AHC mutation, without indels that would impair primer site binding and allele amplification during PCR.

### 2.4 Neural differentiation from iPSCs: Shi cortical protocol

iPSCs were differentiated toward a cortical glutamatergic fate as described previously (Neely et al., 2012; Shi et al., 2012; Telias et al., 2014) with minor modifications. iPSCs were maintained on Matrigel in mTeSR1 and incubated at 37°C in the presence of 5% CO_2_ until they reached approximately 70-80% confluency. iPSC colonies were passaged with Accutase and plated at high density (25×10^4^ cells / cm^2^) in mTeSR1 with ROCK-inhibitor onto Matrigel-coated 12-well plates (day -2). The cells were fed mTeSR1 the following day (day -1), reached confluency, and switched to neural induction media (Shi NIM: Neural maintenance media [NMM] supplemented with 10 μM SB-431542 and 100 nM LDN-193189) on day 0. Neuroepithelial sheets were passaged using dispase on day 10 at a ratio of 1:1 into Matrigel-coated 6-well plates. The following day (day 11), cultures were fed NMM. Upon the appearance of rosettes (around day 16), NMM was supplemented with 20 ng/mL FGF2 for 3 days before being withdrawn. Cells were passaged at day 20 with dispase at a ratio of 1:2-3 onto Matrigel-coated 6-well plates and expanded until day 32 in mixed cortical neural differentiation media (Mixed Cortical NDM supplemented with 10 ng/mL BDNF, 10 ng/mL GDNF, 10 ng/mL IGF-1, and 1 μM dibutyryl cAMP). At day 32, iPSC-derived neurons were dispersed to single cells with Accutase and plated onto destination or maintenance plates for further analysis and culture. Half media changes were performed every 2-3 days as determined by media usage. Neural media recipes and full reagent list are included in the supplemental methods.

### 2.5 Neural differentiation from iPSCs: Maroof GABAergic protocol

iPSCs were differentiated toward a GABAergic cortical interneuron fate as described previously (Maroof et al., 2013) with slight modifications. iPSCs were plated to confluency in a 12-well plate as described above. At d0, media was switched to Maroof NIM (KSR media supplemented with 10 μM SB-431542, 100 nM LDN-193189, and 2 μM XAV939). A titration of the base media from KSR to NMM occurred during days 5-7 of differentiation, with small molecules present until passage at day 10. Differentiating cultures were passaged using dispase on day 10 at a ratio of 1:1 into Matrigel-coated 6-well plates into NMM containing 50 ng/mL sonic hedgehog (SHH) and 1 μM purmorphamine (NMM+PurSHH). Cultures were fed daily with NMM+PurSHH until day 18. Cells were passaged at day 20 with dispase at a ratio of 1:2-3 onto Matrigel-coated 6-well plates and expanded until day 32 in GABAergic neural differentiation media (GABAergic NDM supplemented with 10 ng/mL BDNF, 10 ng/mL GDNF, 200 μM ascorbic acid, and 200 μM dibutyryl cAMP). At day 32, iPSC-derived neurons were dispersed to single cells with Accutase and plated onto destination plates. Half media changes of were performed every 2-3 days as determined by media usage.

### 2.6 iPSC and neural immunostaining

For pluripotency validation in iPSCs and for immunostaining of embryoid body aggregates, cells were washed with PBS and fixed with 4% PFA for 20 minutes at room temperature. Samples were permeabilized with 0.1% TritonX-100 in PBS (PBST) for 5 minutes and blocked with a solution of 2.5% normal goat serum (NGS) in PBST for 1 hour at room temperature. This protocol was modified for staining iPSC-derived neurons to preserve cellular architecture, with neurons fixed in 2% PFA and the blocking solution included 5% NGS. Primary antibodies were applied in the blocking solution overnight at 4°C followed by three PBS washes. Secondary antibodies were applied in blocking solution followed by another PBS wash before nuclear staining with DAPI and mounting on coverslips or plate storage at 4C, if necessary. Low magnification (4-20x) images were collected using an EVOS fluorescent microscope. High magnification (60x) images were acquired using an Andor DU-897 EMCCD camera mounted on a Nikon spinning disk microscope. Confocal images are shown for qualitative assessment of membrane expression of NKA-α3 subunits. Software used for image acquisition and reconstruction included NIS-Elements Viewer (Nikon) and ImageJ (FIJI).

### 2.7 Immunoblotting

Protein was collected from iPSCs at d0 and from neurons at d32 and d60 of differentiation using ice-cold RIPA buffer containing protease and phosphatase inhibitors. Protein was not boiled prior to electrophoresis to avoid aggregation of NKA-α3, a membrane associated protein. Ten micrograms of total protein mixed with 4x Laemmli loading buffer (BioRad) per lane were run on Criterion XT Bis-Tris 4-12% polyacrylamide gels and separated in MOPS buffer at 200V for 60 minutes. Protein was transferred to PVDF membranes at 95V for 65 minutes. Membranes were washed with TBS three times before being stained for total protein with REVERT (Li-Cor). The protein stain was removed, blots were washed three times with TBS, and the membranes were blocked with Odyssey Blocking Buffer for 60 minutes at room temperature. Primary antibodies were applied overnight at 4°C in an antibody incubation solution of TBS + 0.1% Tween-20 (TBST) with 5% BSA. See supplementary antibody Table for more information on usage and dilutions. Blots were washed five times with TBST the following day, and secondary antibodies were applied for two hours at room temperature in antibody incubation solution. Membranes were washed five times with TBST prior to image capture on an Odyssey Scanner. Intensity data was collected using Image Studio Lite. Data was normalized to total protein (see Supplemental Figure 4) with β-Actin shown as a visual aid.

### 2.8 RT qPCR

RNA was collected from iPSCs at d0 and iPSC-derived neurons at d32 and d60 of differentiation (Qiagen RNeasy Kit, QIAshredder) and processed with on-column DNase treatment to remove genomic DNA, followed by long-term storage at −80C. cDNA was generated using the High Capacity cDNA RT Kit (Applied Biosystems, 4368814) and stored at −20°C until further processing. Custom formatted qPCR plates were ordered from Applied Biosystems to profile for relevant NKA subunits with primers and TaqMan probes described in the supplementary reagents table. cDNA samples were prepared using TaqMan PCR Master Mix (Applied Biosystems, 4304437), and data was collected on a QuantStudio 3 qPCR machine. Analysis was performed using averages of technical duplicates for each sample. Calculations were performed using the 2^(− ΔΔC_t_) method, normalizing to GAPDH and then comparing results to expression of *ATP1A1* mRNA in clone IC1 at d0 to assess NKA subunit transcript expression across differentiation timepoints.

### 2.9 MEA culture and flunarizine treatment

For Shi protocol cortical cultures, Axion Cytoview 48-well MEA plates were pretreated with a 0.1% polyethyleneimine (Sigma, P3143) in borate buffer solution for one hour, washed thoroughly with water, and dried overnight the day prior to plating. For Maroof Protocol GABAergic cultures, a thin coat of Matrigel was used instead of PEI as this significantly enhanced adhesion to the electrode recording surface during extended culture periods. Neurons were dispersed with Accutase at d45 of differentiation and plated at a concentration of 3×10^5^ cells/well into MEA plates in 50 μL of culture media containing 10 μg/mL laminin (Sigma, 11243217001). One hour after plating, the well was gently flooded with 300 μL culture media (NDM). From MEA plating onward, culture media included B27 Plus (Gibco) instead of standard B27 to promote neuronal activity. MEA cultures were fed on d48 with a half-media change and visualized under light microscope to ensure coverage of the electrode surface. At and after d51, half-media changes with media including 1 μg/mL laminin were performed every other day, including 24 hours before recording sessions. From d59 onward, media included 0.1% DMSO vehicle or 100 nM flunarizine (Sigma, F8257) for treatment groups.

### 2.10 MEA recordings and heat stress modeling of trigger-induced phenotypes

MEA recordings were performed every 4 days beginning at d52 on an Axion Maestro Pro MEA system. The recording chamber was preheated to 37°C with CO_2_ concentration set to 5%. Spontaneous neural activity was recorded with a sampling frequency of 12.5 kHz and a digital bandpass filter from 200 Hz to 3 kHz. For validation that these settings were capturing neural signals, a dose curve with TTX (Abcam, ab120055) was performed on an initial MEA plate of iPSC-derived neurons not included in the main differentiation analysis presented here. Plates were pre-incubated for 10 minutes within the MEA system before a 10-minute baseline recording was initiated. At the end of this recording period, the temperature was rapidly increased to 40°C and a new 10-minute “heat stress” recording period started once the temperature reached 39.5°C. When this recording period concluded, the temperature was reset to 37°C and a 10-minute “recovery” period began when the chamber temperature fell to 37.5°C. Data were analyzed using AxIS 2.0 Software (Axion Biosystems) by replaying recordings in offline neural spontaneous mode. For each individual recording, activity metrics from four wells of the same clone and treatment condition were summed to control for internal variation between wells and days. Pooled values of less than 50 total spikes (5 spikes/minute) in a baseline period resulted in the exclusion all three heat conditions (baseline, heat stress, and recovery) for that recording session in ratiometric analyses. One set of heat stress recording data was excluded from a single differentiation and day for clone UC1 given abnormally high heat stress and recovery period activity ratios, as determined by Grubbs’ test for outliers.

### 2.11 Data analysis

Statistical analysis was performed using Prism 8 (GraphPad). Two iPSC clones for each genotype (unrelated *ATP1A3*^+/+^, isogenic *ATP1A3*^+/+^, AHC *ATP1A3*^+/E815K^) over 5 differentiation runs are reported for all iPSC-derived neuron data sets. For analysis of multiple comparisons where n=10, a Bonferroni-correct threshold p-value of < 0.005 was considered statistically significant. Data are presented as mean ± SEM. Relevant statistical tests and sample sizes are indicated in figure legends.

## 3. Results

### 3.1 Generation of E815K patient-specific iPSCs and isogenic wildtype controls

We employed episomal plasmid reprogramming methods to generate iPSC lines from fibroblasts of an AHC patient harboring a heterozygous missense mutation in exon 18 of *ATP1A3* (c.2443G>A, p.E815K; AHC *ATP1A3*^+/E815K^; lines AHC-20 and AHC-24). Clones from two unrelated individuals were also used as wildtype controls (unrelated *ATP1A3^+/+^;* lines UC1 and UC2). iPSC clone pluripotency was confirmed by immunostaining for common pluripotency markers, successful germline marker identification in trilineage differentiation assays, and the presence of a normal karyotype (**Supplemental Figure 1**). PCR assays were performed to ensure that episomal plasmids used in reprogramming had not integrated into the host genome. CRISPR/Cas9 genome editing was utilized to create verified isogenic iPSC wildtype clones (isogenic *ATP1A3*^+/+^; lines IC1 and IC2) (**Supplemental Figure 2**). Two clones of each genotype (unrelated *ATP1A3^+/+^*, isogenic *ATP1A3*^+/+^, AHC *ATP1A3*^+/E815K^) were used in an iPSC-derived neuronal model of disease to discover and analyze *in vitro* phenotypes caused by the AHC E815K mutation (**Figure 1A**). Sequencing of the *ATP1A3* region surrounding the mutation of interest was performed to confirm the expected mutation status of each line (**Figure 1B**) prior to the beginning of differentiation runs.

**Figure 1:**
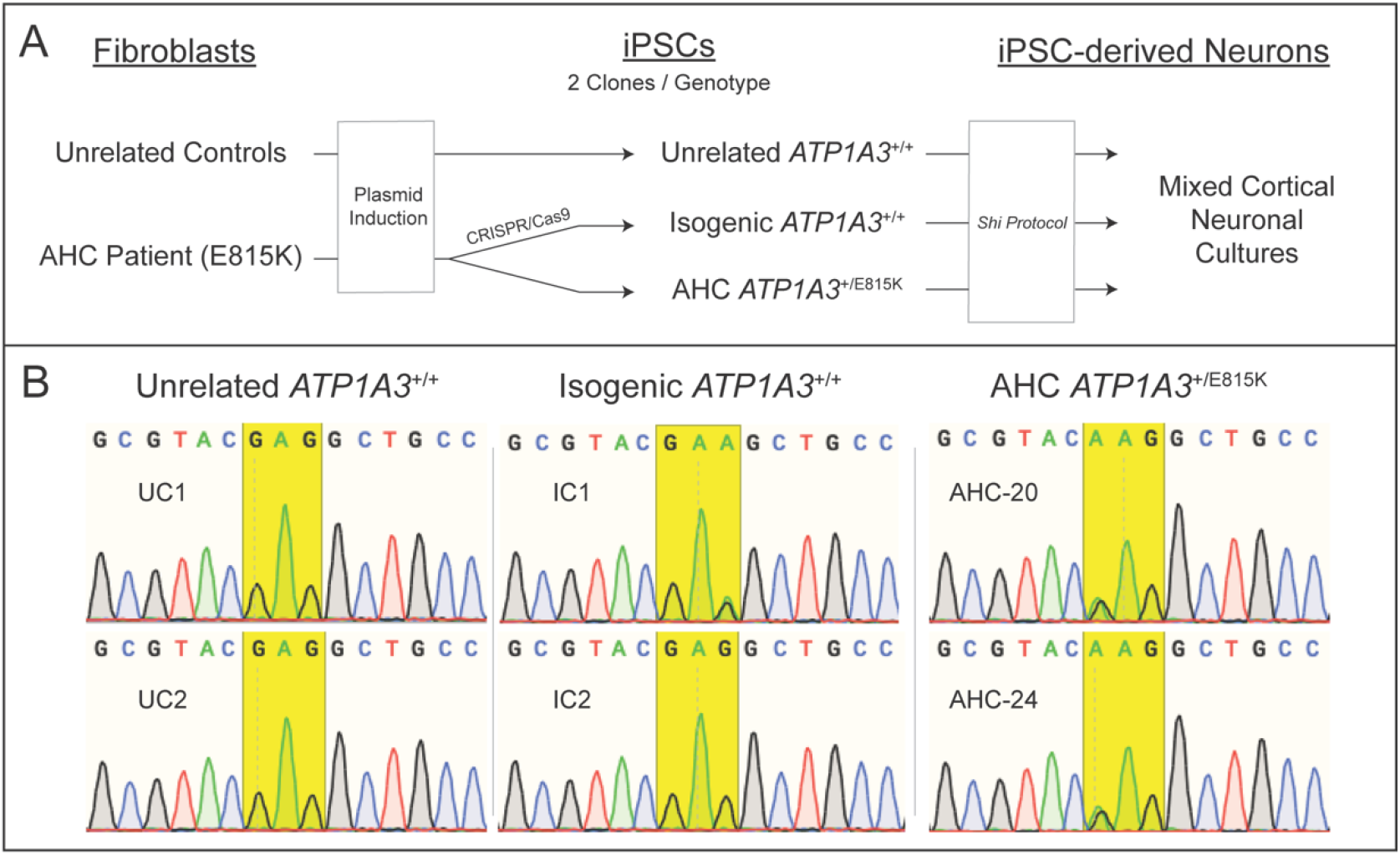
Experimental approach and sequencing of iPSC clones used in this study. (A) Fibroblasts were collected by punch skin biopsy from two unrelated control volunteers (*ATP1A3*^+/+^) and from a patient with AHC harboring the *ATP1A3*^+/E815K^ genotype. Cultured fibroblasts were induced to pluripotency using episomal transduction methods. Isogenic wildtype clones (*ATP1A3*^+/+^) were generated from a patient line to be used as an additional control group. Two clones of each genotype (unrelated, isogenic, AHC) were differentiated to mixed cortical neuronal cultures to be analyzed in this study. (B) Sequencing of PCR products for the region surrounding the mutation site of interest in exon 18 of *ATP1A3* show the wildtype status in unrelated control (UC) iPSC clones, corrected sequences in isogenic control (IC) clones, and the pathogenic c.2443G>A heterozygous status in AHC clones. The yellow box highlights the sequence encoding the 815^th^ residue. Intended alterations of the PAM-site to prevent repetitive Cas9 cutting events result in additional synonymous changes in isogenic clones. As unique gRNAs were used to create separate isogenic iPSC clones, a different synonymous PAM-site change exists further upstream in clone IC2 not shown in this sequencing read. See Supplemental Figures 1 and 2 for validation data, expanded isogenic clone chromatographs, and genetic editing strategy.

### 3.2 AHC patient iPSCs successfully differentiate into neurons

We used a previously published protocol for two-dimensional dual-SMAD inhibition based neural differentiation (Shi et al., 2012) to generate early excitatory neuronal cultures by mimicking cortical plate formation (**Figure 2A**). Cultures resulting from this differentiation protocol are mainly comprised of glutamatergic neurons. NKA-α3 subunit protein was detected by immunostaining in iPSC-derived neurons and was clearly expressed in cells robustly positive for neural markers β-III-Tubulin and MAP2 (**Figure 2B**). These cultures eventually form “mixed cortical” populations including small subsets of GABAergic neurons. Substantial numbers of glial cells are not generated in this protocol until after the time periods analyzed in this study (d80+). Our experimental approach was modified to include standardized passage dates, morphogen treatment windows, and data collection timepoints throughout differentiation (**Figure 2C**). To minimize the influence of inherent variation of iPSC-derived neuronal cultures on our results, five independent differentiations were conducted and analyzed.

**Figure 2:**
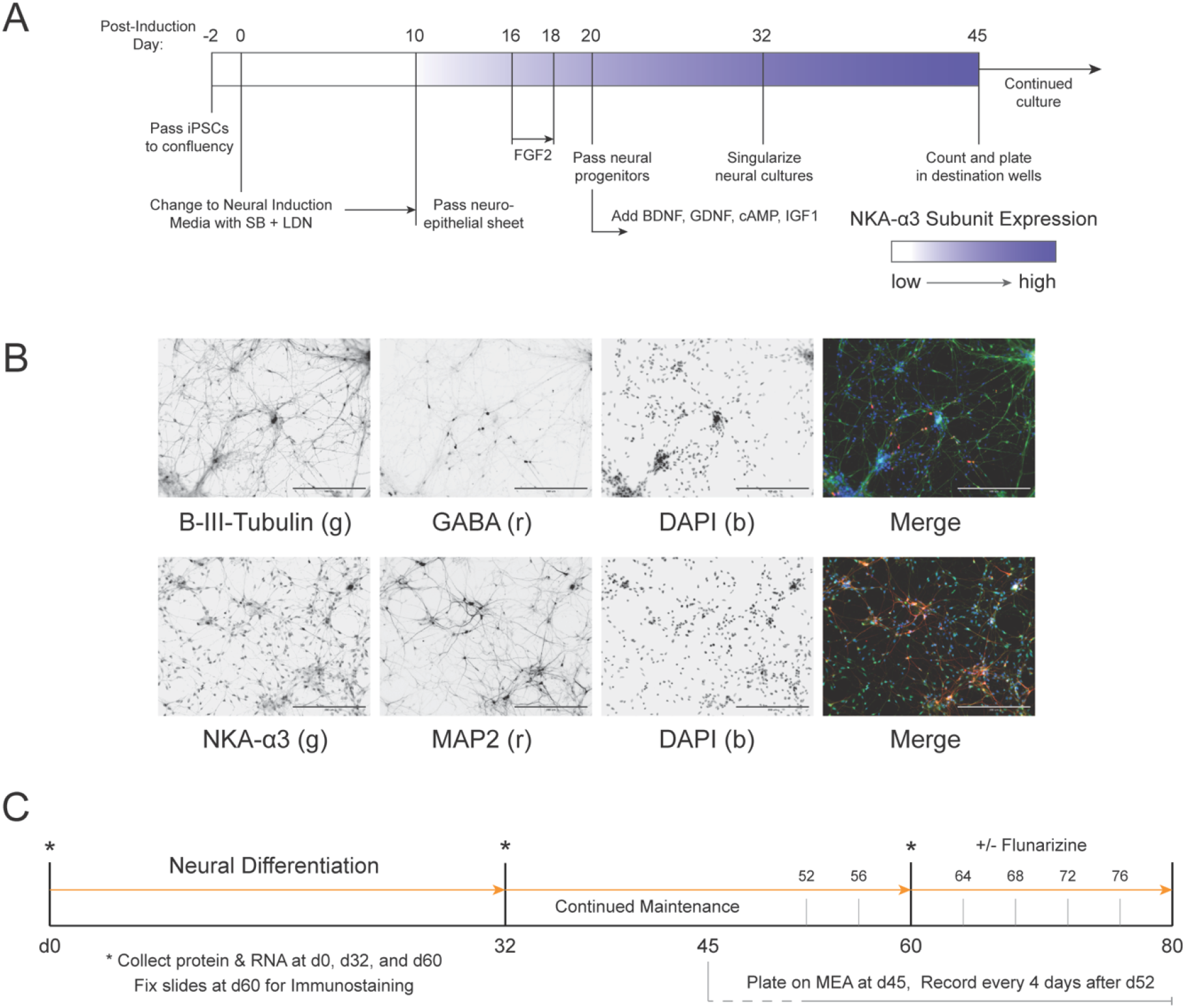
Neural differentiation protocol, outcomes, and experimental timeline. (A) A slightly modified version of a previously published differentiation protocol (Shi et al., 2012) was used to generate the disease model. This protocol uses dual-SMAD inhibition based neural induction with SB-431542 and LDN-193189 to generate a neuroepithelial sheet by d10 of differentiation. FGF2 treatment and withdrawal upon the appearance of neural rosettes results in expansion of neural precursors. Growth and maturation factors were added at d20 of induction and cultures were dispersed to individual neurons at d32. After continued expansion, neurons were plated on destination wells (MEA or chamber slides) at d45. (B) Immunostaining of AHC cultures fixed at d60 shows the expression of neural markers including β-III-Tubulin and MAP2. Cultures at this stage begin generating small populations of GABAergic lineages in addition to the dominant glutamatergic populations described in the original publication. Neurons generated in these cultures stain positive for the NKA-α3 subunit, with no obvious differences between genotypes or intracellular aggregations observed. DAPI shown as a nuclear stain; scale bars = 200 μm. (C) The experimental timeline for this study included three collection timepoints for protein and RNA (*; d0, d32, d60). Neurons were plated onto microelectrode array (MEA) plates at d45 and recordings began at d52 through d80 at 4-day intervals. From d60-d80, 100 nM flunarizine or vehicle (DMSO) was added to the MEA feeding media.

### 3.3 iPSC-derived neurons express NKA-α3 subunit protein early during in vitro neurodevelopment

Evidence from heterologous overexpression studies suggest that while some *ATP1A3* mutations have the potential to cause trafficking abnormalities, no known AHC causing mutations result in altered membrane expression of the NKA-α3 subunit (de Carvalho Aguiar et al., 2004; Heinzen et al., 2014). We hypothesized that our *in vitro* human stem cell derived model would show no changes in NKA-α3 protein expression and localization. Immunostaining for NKA-α3 did not reveal any obvious localization differences or intracellular aggregation between AHC and control iPSC-derived neurons (**Figure 2B, Supplemental Figure 3**). Recent studies have explored the possibility of competitive relationships between NKA-α1 and NKA-α3 in phenotypic expression of disease variants (Arystarkhova et al., 2019). We therefore performed immunoblotting for these proteins to analyze NKA-α1 and NKA-α3 expression relationships in control and disease groups throughout the course of *in vitro* neural differentiation (**Figure 3A**). No significant differences in differentiation toward a neuronal fate by β-III-Tubulin expression were noted between controls and disease at d32 or d60 of differentiation (**Figure 3B**). NKA-α1 expression dropped during early neural development at d32 then rebounded by d60 of differentiation, with no differences between control lines and AHC patient lines (**Figure 3C**). The neuronal specific NKA-α3 subunit, as expected, was absent in iPSCs before rising substantially during neural differentiation. No differences in NKA-α3 subunit expression existed at d32 between disease and control neurons, while at d60 α3 expression was significantly elevated in *ATP1A3*^+/E815K^ mixed cortical cultures compared to unrelated controls (p = 0.0047). However, no significant difference was noted compared to isogenic controls (p = 0.7257), highlighting the importance of incorporating isogenic lines in experimental disease modeling when possible (**Figure 3D**). Overall, data trends support nonsignificant changes in neural differentiation outcomes and NKA-α subunit protein expression in *ATP1A3*^+/E815K^ disease lines compared to control lines.

**Figure 3:**
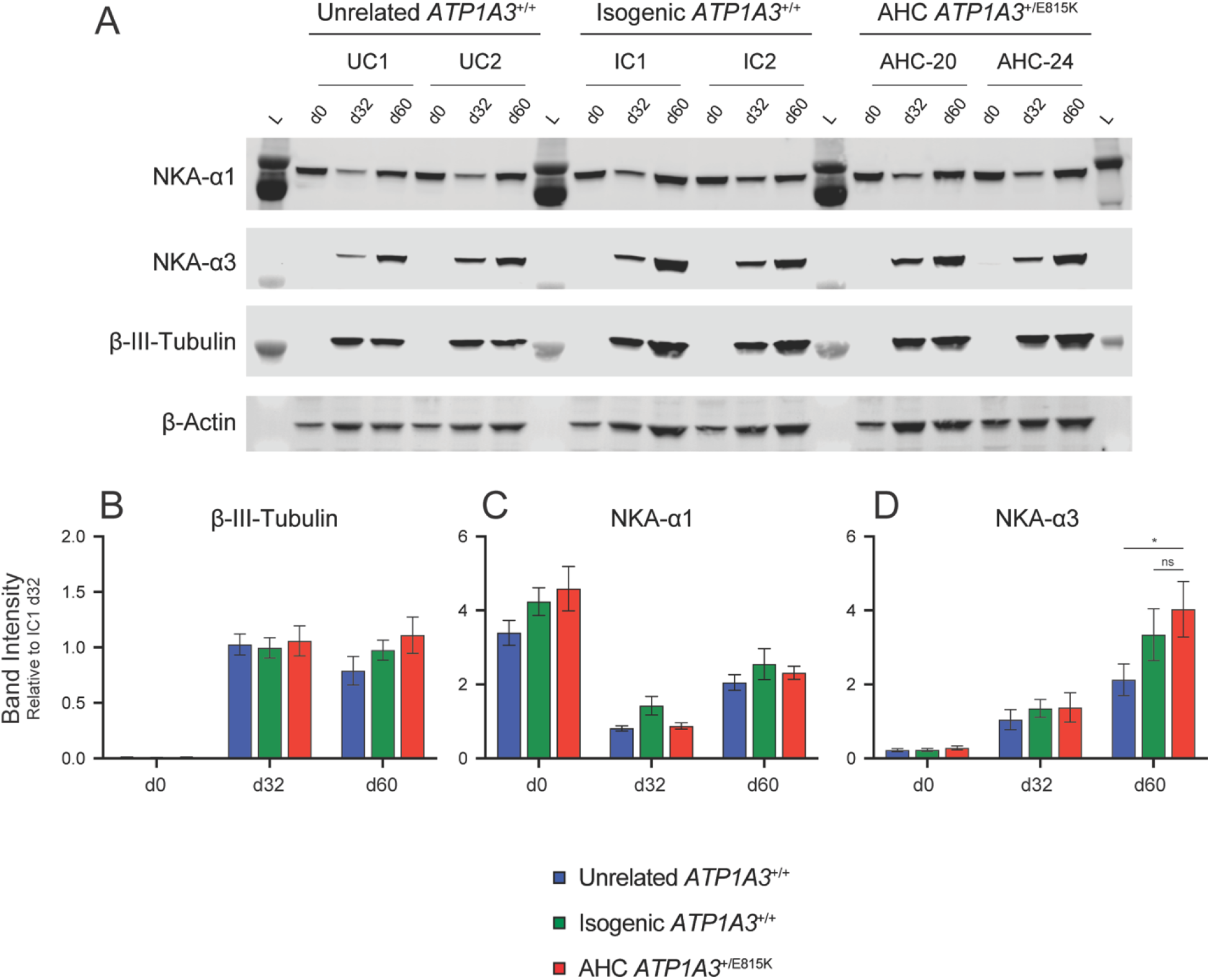
AHC and control iPSC-derived neurons express similar levels of NKA-α3 subunit protein during *in vitro* neurodevelopment. (A) Representative immunoblot of protein collected from mixed cortical neuronal cultures and probed for NKA-α1, NKA-α3, and β-III-Tubulin. While β-Actin is shown as a visual guide for protein loading, intensity data was normalized to total protein by REVERT staining show in the supplementary data. (B) β-III-Tubulin expression increases as expected during iPSC-derived neuronal induction. No significant differences in induction efficiency by this measure are noted on immunoblot between genotypes. (C) NKA-α1 expression dropped during early neurodevelopment at d32 then increased by d60 in a consistent pattern with no significant differences between genotypes. (D) NKA-α3 expression was very low to nondetectable in iPSCs at d0 and increased drastically by d32 and d60 of differentiation. AHC lines expressed significantly more NKA-α3 by immunoblot compared to unrelated control lines (p = 0.0047), while this difference was not observed in isogenic corrected and AHC lines (p = 0.7257). Graphs display quantified intensity data relative to clone IC1 d32 intensity. n = 10 (5 differentiations, 2 clones per genotype). 2-way ANOVA with Bonferroni’s multiple comparisons test; significant findings notated by *, p < 0.005; ns = not significant (p > 0.005).

### 3.4 AHC neurons display increased NKA-α3 subunit transcripts

While no statistically significant differences were noted for NKA-α1 and NKA-α3 subunit protein expression between isogenic control and AHC lines during neural differentiation, this does not rule out differences in transcriptional regulation of these or other related NKA subunits. To assess whether transcriptional compensation of these subunits existed in our disease model, we profiled NKA-subunit transcription of iPSC-derived neurons at d0, d32, and d60 of differentiation. *ATP1A4* transcript levels were not assessed as this gene product has been shown to be expressed only in a testes-specific pattern, with no expression observed in our model during preliminary studies. iPSCs expressed predominantly *ATP1A1* and *ATP1B3* transcripts (**Figure 4A**), with some expression of *ATP1A2* and other β-subunit transcripts; these patterns were consistent between genotypes. While there was a significant elevation in unrelated control cells of *ATP1A2* mRNA compared to the disease and isogenic control groups, there were no differences in any other transcripts. The NKA-subunit expression profile shifts toward an *ATP1A3 and ATP1B1* dominant pattern in iPSC-derived neurons. Intriguingly, there was a significantly greater level of *ATP1A3* transcript expression in disease neurons compared with control groups at both neural collection timepoints (AHC-to-isogenic: d32 p = 0.0015; d60 p = 0.0005). These patterns remain consistent whether comparing AHC transcript levels to unrelated or isogenic control data (**Figure 4B-C**). Later significant elevation of *ATP1B1* transcripts was also noted, suggesting the possibility of a lagging compensation of this interacting subunit. These data suggest the existence of a feedback mechanism in iPSC-derived neural cultures resulting in elevated expression of neural specific NKA-subunit transcripts in the presence of a disease causing AHC mutation and associated cellular abnormalities.

**Figure 4:**
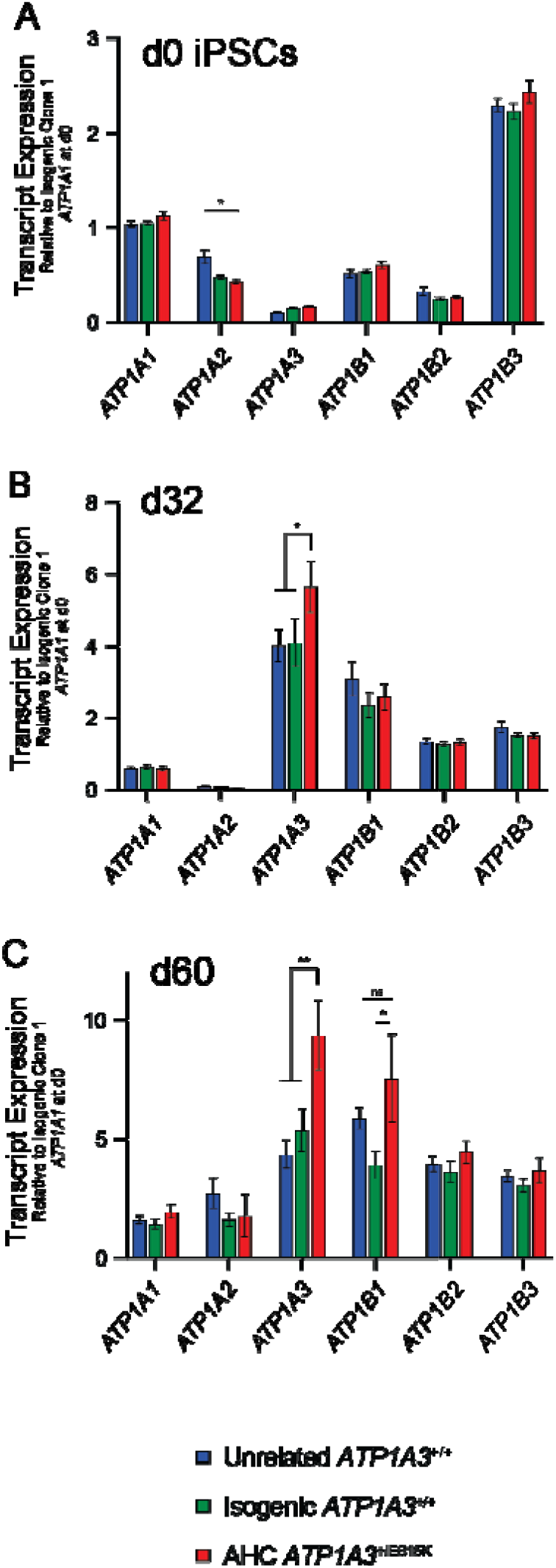
The transcriptional profile of NKA-subunits is altered in AHC patient iPSC-derived neurons. (A) The NKA transcriptional profile of iPSCs is dominated by *ATP1A1* and *ATP1B3* expression. While *ATP1A2* transcripts are elevated in unrelated *ATP1A3*^+/+^ iPSCs compared to AHC lines (p = 0.0010), there is no difference noted between AHC and isogenic corrected iPSCs (p > 0.9999). (B) By d32 of *in vitro* neuronal differentiation, the NKA-subunit transcription pattern shifts toward *ATP1A3* and *ATP1B1* expression. There was a significant elevation at this timepoint in *ATP1A3* transcript levels between AHC and both isogenic (p = 0.0015) and unrelated (p = 0.0008) control neurons. (C) At d60, elevated production of *ATP1A3* mRNA was maintained (AHC-to-isogenic p = 0.0005; AHC-to-unrelated p < 0.0001), and a lagging elevation of *ATP1B1* mRNA was noted in AHC groups compared to isogenic corrected neurons (p = 0.0018). Graphs display transcript expression relative to clone IC1 *ATP1A1* transcripts at d0. Statistical conclusions are maintained if normalizing to *ATP1A1* expression values for each individual clone at d0. n = 10 (5 differentiations, 2 clones per genotype). 2-way ANOVA with Bonferroni’s multiple comparisons test; significant findings notated by: * p < 0.005, ** p < 0.001; ns = not significant (p > 0.005).

### 3.5 iPSC-derived neurons are electrically active on MEA analysis

To assess if AHC patient derived neurons manifest differences in neural activity on a population level, we interrogated iPSC-derived neurons using microelectrode array (MEA) recordings. Activity was first recorded at d52 and continued every 4 days until cultures were terminated at d80. Coculture with glial cells was not performed in an attempt to isolate changes specific to this differentiation protocol. MEA plates were visually observed after plating to ensure coverage of the recording area (**Supplemental Figure 5A**), and neural spike recording parameters are supported by complete activity inhibition by TTX (**Supplemental Figure 5B**). Neurons from this mixed cortical differentiation protocol were active on MEA analysis, and neural activity of iPSC-derived cultures continued to increase over the course of d52-to-d80 MEA analyses (**Figure 5A**). While AHC lines trended towards lower activity compared to both control groups at multiple timepoints, no statistically significant phenotypes were noted. Chronic treatment with flunarizine, a drug commonly used to reduce the severity and frequency of AHC episodes was assessed to identify any impact of treatment in disease versus control lines (**Figure 5B**). A dose of 100 nM was chosen based on an initial dose-response experiment where concentrations at or above 500 nM caused significant loss of neuronal activity in all genotypes (**Supplemental Figure 5C**). Flunarizine-treated groups displayed similar dynamics to vehicle-treated groups, although flunarizine generally prevented the time-dependent activity increase in neuronal cultures without significant differences between control and E815K mutant groups. While flunarizine treatment decreased neural activity levels in both isogenic control and AHC disease lines, unrelated control groups did not have a large activity change in the presence of this drug further stressing the importance of including isogenic cells in study designs (**Supplemental Figure 5D**).

**Figure 5:**
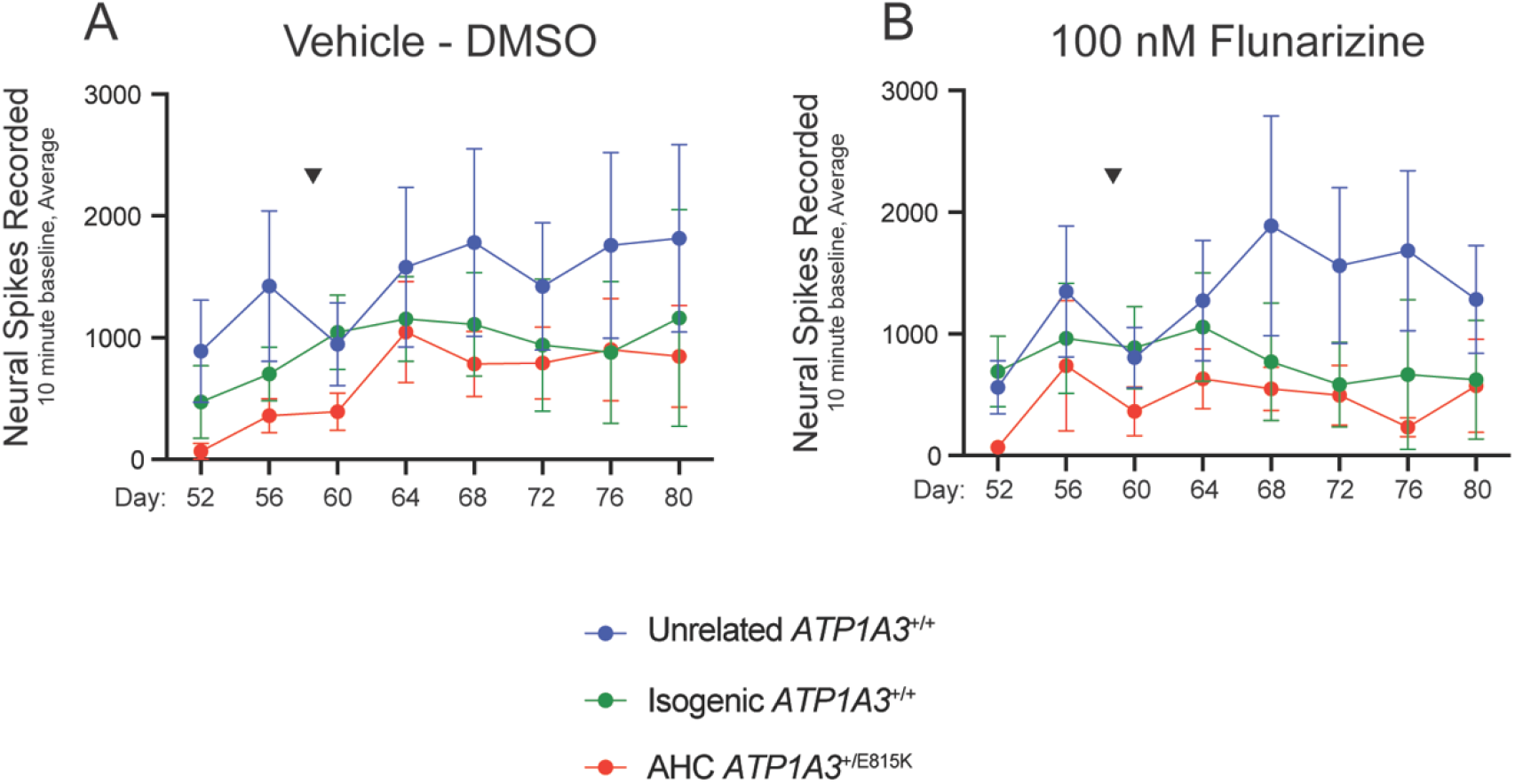
iPSC-derived neurons are electrically active by microelectrode array (MEA) analysis. Mixed cortical neuronal cultures were chronically treated beginning at d59 (arrowheads) with either DMSO (A) or 100 nM flunarizine (B). Regardless of treatment, baseline recordings of spontaneous neural activity show electrical activity and maturation over the course of the 28-day recording window. Electrical maturation was limited following chronic treatment with flunarizine, particularly in AHC and isogenic corrected neurons as quantified in the supplement. While unrelated wildtype and isogenic corrected neurons trend toward increased overall activity compared to AHC neurons, particularly at the beginning of the recording period, variability in spontaneous activity is too high to allow for statistically significant conclusions. n = 10 (5 differentiations, 2 clones per genotype) for each recording day; data collected on Axion Maestro Pro using spontaneous neural activity acquisition settings.

### 3.6 Trigger-induced neural hyperactivity in AHC cultures

Trigger-induced hemiplegic spells are a pathognomonic aspect of AHC and thus a possible insight into mechanistic underpinnings of the disease and related disorders. Described triggers range from fever to exposure to water, ingestion of specific foods, and infection, among others. Animal models of disease have replicated stress-induced phenotype exacerbation, including vestibular stimulation or chronic restraint causing hemiplegia and dystonia in mouse models and heat-induced paralysis in *Drosophila* models of *ATP1A3* disease (Helseth et al., 2018; Holm and Lykke-Hartmann, 2016; Isaksen et al., 2017; Palladino et al., 2003; Sugimoto et al., 2014). To model triggered phenotypes in an iPSC-derived neuronal system, cultures on MEA plates were exposed to a heat stress protocol beginning at d60 of differentiation. Separate recordings of 10-minutes in duration were taken at baseline (37°C), with heat stress (40°C), and during recovery (37°C) periods in the presence (**Figure 6A**) or absence (**Figure 6B**) of chronic flunarizine. When normalizing to baseline activity for each clone per recording session, heat stress resulted in a slight reduction of activity that was insignificantly different between genotypes. However, a robust rebound hyperactivity phenotype was observed in AHC neurons compared to both the unrelated and isogenic control lines (p < 0.0001 for both comparisons). Chronic flunarizine treatment did not affect this phenotype compared to isogenic controls (AHC-to-isogenic p < 0.0001; AHC-to-unrelated p = 0.0119). However, when analyzed by individual recording day (**Supplemental Figure 6**), flunarizine treated cultures had fewer days with significant differences (2/6) compared to vehicle treatment (4/6).

**Figure 6:**
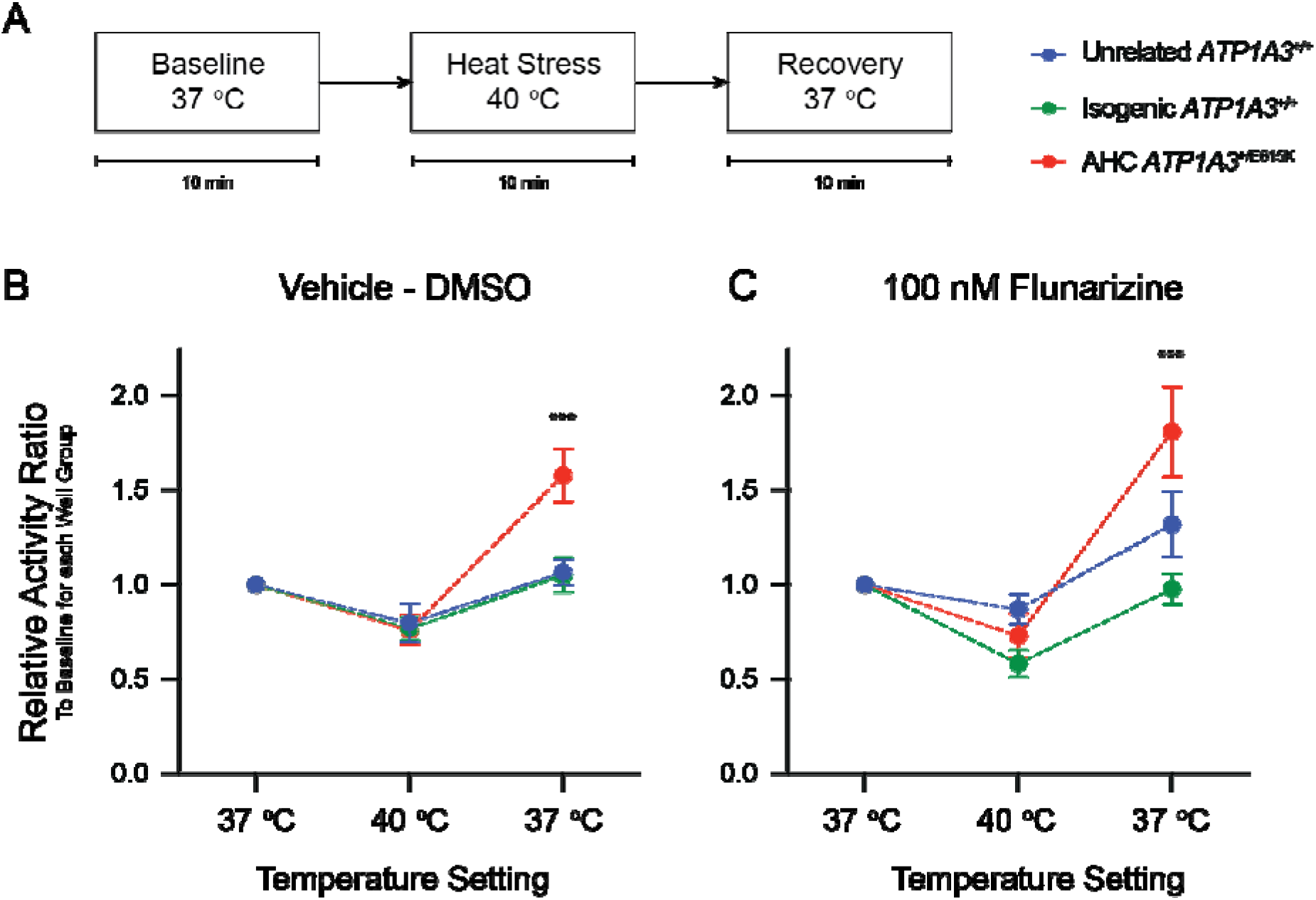
AHC neurons display a stress-induced hyperactivity phenotype following exposure to increased temperature. To assess if this *in vitro* disease model recapitulated features of trigger-induced episodes in AHC patients, a paradigm of increased temperature and recovery (A) was designed for use on the MEA recording device. Spontaneous neural activity in cultures was recorded at a baseline temperature of 37°C, after which the environmental temperature was increased to 40°C followed by a recovery period at 37°C. Each recording period had a duration of 10 minutes. Activity of well groups for each clone and treatment condition was normalized to the baseline period to eliminate inherent variability in activity across different recording days and differentiations. (B) In the presence of elevated temperature, spontaneous neural activity dropped to approximately 75-85% of baseline activity across all genotypes. During the 10-minute recovery period, a relative hyperactivity phenotype was noted in the AHC neurons compared both unrelated wildtype and isogenic correct control neurons (recovery period AHC-to-isogenic p < 0.0001; AHC-to-unrelated p = 0.0001). (C) Recovery phase hyperactivity in AHC neurons is unchanged following chronic 100 nM flunarizine treatment. (recovery period AHC-to-isogenic p < 0.0001; AHC-to-unrelated p = 0.0119) Baseline recording groups with activity values of <50 spikes (5 spikes / minute) resulted in the exclusion of that heat stress data set for the recording day. Heat stress protocol was repeated across 6 recording days (d60, d64, d68, d72, d76, and d80), with each subsequent recording being treated as a technical replicate for that differentiation. n = 10 (5 differentiations, 2 clones per genotype). Data per recording day is displayed in the supplemental data. 2-way ANOVA with Bonferroni’s multiple comparisons test; significant findings notated by: *** AHC-to-isogenic p < 0.0001.

### 3.7 Findings are broadly replicated using a GABAergic differentiation protocol

A distinct protocol that drives iPSCs toward cortical GABAergic interneuron fates was used to analyze if our findings were specific to the original glutamatergic predominant cortical protocol or representative of a more generalized feature of AHC patient stem cell derived neurons. A previously published GABAergic differentiation protocol (Maroof et al., 2013) was slightly modified to synchronize passage timepoints with the mixed cortical protocol, using Wnt-antagonism and Sonic Hedgehog (SHH) treatment to generate larger populations of ventral forebrain lineages in culture (**Supplemental Figure 7A-B**). While early neural cultures at d32 were consistently pure using this protocol, contaminating non-neuronal populations became obvious by d60 as demonstrated by loss of β-III-Tubulin staining by immunoblotting (**Supplemental Figure 7C**). At d32, similar profiles of NKA-α1 and NKA-α3 protein expression existed compared to the mixed cortical differentiation protocol (**Supplemental Figure 7D-E**). Importantly, a significant elevation of *ATP1A3* transcript expression in disease over control groups was again noted at d32 (p = 0.0003) that deteriorated with loss of culture purity by d60 (**Supplemental Figure 7F-G**). Transcriptional differences were again consistent when comparing AHC transcript levels to either unrelated or isogenic controls at d32. Due to culture impurity at d60+, MEA recordings were unable to record sufficient levels of activity during for relevant analysis of GABAergic neuronal cultures.

## 4. Discussion

Alternating hemiplegia of childhood is the prototypical member of an expanding spectrum of disorders attributable to mutations in *ATP1A3* (Brashear et al., 2018; Rosewich et al., 2012; Sweney et al., 2015; Yano et al., 2017). *ATP1A3* spectrum disorders are almost exclusively caused by heterozygous *de novo* missense mutations in this essential gene that encodes the neuronal-specific catalytic subunit of the Na,K-ATPase ion transporter. These distinct diseases often share symptomatic profiles but have unique diagnostic criteria and strong genotype-to-phenotype correlations (Heinzen et al., 2014; Rosewich et al., 2014). While a singular molecular mechanism is unlikely to be responsible for all of these phenotypes, uncovering the biological basis of AHC may yield important information about the others.

The majority of AHC patients harbor one of three unique heterozygous missense mutations: D801N, E815K, or G947R (Ishii et al., 2013; Viollet et al., 2015). These mutations cluster around ion binding pockets within the transmembrane core and result in decreased pump function (Koenderink et al., 2003; Sweadner et al., 2019; Weigand et al., 2014). Heterologous expression studies demonstrated that AHC mutations result in impaired ion transport including loss of proton transport availability particularly in the case of E815K mutations (Li et al., 2015; Weigand et al., 2014). Mouse models of *ATP1A3* disease often recapitulate some symptoms experienced by AHC patients, especially following stress. For instance, a previously published E815K mouse model demonstrated increased hemiplegic spells following vestibular stimulation-induced stress compared to control animals (Helseth et al., 2018). Electrophysiological recordings in mouse hippocampal slices in D801N and I810N variant models of AHC have also shown hyperexcitable responses to electrical stimulation or seizures (Clapcote et al., 2009; Hunanyan et al., 2015; Hunanyan et al., 2018), along with studies reporting neural hyperexcitability in the presence of pharmacologically inhibited NKA ion transport (Vaillend et al., 2002). Other studies in the field have demonstrated that these electrophysiological aberrations may be related to intracellular increases in sodium concentration in both the presence of AHC causing mutations and with ion transport inhibition after treatment with pump antagonists (Pivovarov et al., 2018; Tiziano, 2019; Toustrup-Jensen et al., 2014). In collaboration, we recently used iPSC-derived neurons with the G947R mutation to investigate fundamental cellular defects and observed impaired ion pump activity and depolarized resting membrane potentials in AHC mutant neurons (Simmons et al., 2018).

In this study, we chose to study the E815K mutation as this common AHC variant is often associated with the most severe patient phenotype (Panagiotakaki et al., 2015; Sasaki et al., 2014). Using patient-specific iPSCs along with isogenic and unrelated controls, we created an iPSC-derived neuronal model of AHC to investigate possible mechanisms underlying disease pathogenesis. We found elevated expression of *ATP1A3* transcripts in AHC lines compared to controls without significant changes in protein expression. Previous studies have shown that the complex transcriptional regulation of NKA-subunits includes pump inhibition-induced increases in NKA-subunit transcripts across various cell types (Li and Langhans, 2015). Interestingly, reports have implicated elevated intracellular sodium and low intracellular potassium concentrations as transcriptional regulators of NKA-subunit expression patterns (Johar et al., 2014; Vinciguerra et al., 2003; Wang et al., 2007). Our findings identify alterations in transcriptional control as a new phenomenon in *ATP1A3* variant disease, one that may have implications in the development of future therapies. No significant difference in NKA-subunit protein expression was consistently seen across *in vitro* neurodevelopment in this study, as may be expected. This may be due to insensitivity of the available antibodies to detect such changes with immunoblotting or may reflect lower stability of the mutant protein. Higher resolution immunostaining studies may result in more detailed conclusions about protein expression, stability, and subcellular localization in AHC disease models. Prior studies have shown that transcript levels and protein expression are related but not necessarily correlative for all NKA subunits (Clifford and Kaplan, 2009; Devarajan et al., 1992). This is the first report of compensatory transcription of NKA-subunit mRNA in AHC, which has implications for the experimental design and early implementation of genomic therapies targeting mutant or wildtype transcripts, even in the absence of protein-level aberrations. Other AHC-causing *ATP1A3* mutations should be analyzed to validate this observation.

Microelectrode array analyses demonstrated that in iPSC-derived neuronal cultures, *ATP1A3*^+/E815K^ cells trended towards less overall activity than their wildtype equivalents early in the recording period, with stabilization comparable to isogenic control levels over time. This protocol did not yield activity levels representing synchronous network bursting, limiting our ability to perform more complex MEA analysis. However, our approach allowed us to isolate genotype specific changes in activity without necessitating further modifications to the prolonged differentiation protocols. The novel induction of cellular stress by elevated temperature revealed a clear hyperactivity phenotype following heat stress in *ATP1A3^+/^*^E815K^ lines compared to controls. At this time, we do not know if this phenotype is related to pathological neurodevelopment or altered electrophysiological properties associated with a mutant NKA-α3 subunit. Furthermore, the correlation of this phenotype with epilepsy or non-epileptic symptoms of AHC was not determined. The phenomenon of stress-induced triggers is a shared phenotype of AHC with other *ATP1A3* mutant diseases, and has been recapitulated in other disease models including heat stress induced paralysis in a *Drosophila* model (Helseth et al., 2018; Holm and Lykke-Hartmann, 2016; Isaksen et al., 2017; Palladino et al., 2003; Sugimoto et al., 2014). Intriguingly, another study has identified a temperature-sensitive ion leakage phenomenon in a closely related Type II P-Type ATPase family member, after identifying an uncoordination phenotype in *Drosophila* harboring mutations in SERCA (Kaneko et al., 2014). Moving forward, it will be of great interest to test outcomes of stress modeling using newer lentivirally-induced neuron protocols that can generate more consistent network activity (Yang et al., 2017; Zhang et al., 2013).

Flunarizine is one of the most commonly used drugs in AHC to prevent triggered episodes, although patients do not consistently respond to treatment (Panagiotakaki et al., 2015; Pisciotta et al., 2017; Sasaki et al., 2001). Chronic *in vitro* treatment with flunarizine did not have an impact on this stress-triggered phenotype in our study. However, flunarizine may be exerting any influence on other neuronal lineages that we have not addressed. Previous studies have implicated altered GABAergic inhibition as a possible pathological driver of disease (Bottger et al., 2011; Ikeda et al., 2013). We did not observe differences between early glutamatergic and GABAergic neural differentiation protocols, although the failure of GABAergic cultures to maintain neuronal purity and electrical activity on MEA analysis limited our analysis. While NKA signaling has been shown to be important for dendritic growth along with synaptic coordination and maturation, the potential impact on cellular signaling resulting from pump inhibition relative to AHC mutations in human iPSC-derived neurons remains understudied (Aperia et al., 2016; Desfrere L, 2012; Reinhard et al., 2013). Structural modeling has shown similarities between common AHC mutation impacts on ion binding and passage within the Na,K-ATPase catalytic subunit, suggesting a potential broader validity of our findings (Sweadner et al., 2019).

The inclusion of isogenic wildtype controls and the replication of findings in two separate differentiation protocols in this study certainly increased our confidence in the robustness of our findings of both transcriptional changes and heat induced hyperactivity (Germain and Testa, 2017). Our model system can be used in the future to test important questions existing in the *ATP1A3* field regarding disease pathogenesis arising from either haploinsufficiency or dominant negative mechanisms. Furthermore, researchers in the field working on potential new therapeutics, whether small molecule or genetic therapies, can use these findings, model system, and assay techniques to test various future hypotheses on treatment efficacy.

In summary, our iPSC-derived model of AHC demonstrated that neurons from the most phenotypically severe AHC mutation show elevated expression of *ATP1A3* transcripts during *in vitro* neurodevelopment compared to both isogenic corrected and unrelated wildtype controls. Neurons derived from these AHC patient iPSCs form electrically active populations upon MEA analysis similar to controls and display hyperexcitability following exposure to a trigger of elevated temperature. The commonly used medication flunarizine did not rescue this phenotype. Our results provide new evidence for phenotypes in AHC neurons harboring the *ATP1A3*^+/E815K^ genotype and describe a platform for future mechanistic discovery and possibly therapeutic screening.

## Abbreviations

AHC: Alternating hemiplegia of childhood
CAPOS: Cerebellar ataxia, areflexia, pes cavus, optic nerve atrophy, and sensorineural hearing loss
EIEE: Early infantile epileptic encephalopathy
FIPWE: Fever-induced paroxysmal weakness and encephalopathy
MEA: Microelectrode array
NKA: Na^+^, K^+^-ATPase
RDP: Rapid onset dystonia-parkinsonism
RECA: Relapsing encephalopathy with cerebellar ataxia

## Acknowledgements

This research was supported by funding from the Alternating Hemiplegia of Childhood Foundation (K.C.E. and A.L.G.). Additional funding was also provided by Association Française de l’Hémiplégie Alternante. J.P.S. was supported by NIGMS of the National Institutes of Health under award number T32GM007347. We appreciate the helpful comments on this manuscript from members of the Ess laboratory, the assistance of Kevin Lazenby in processing MEA data, confocal microscopy assistance from Mary Chalkley, immunostaining assistance from Brittany P. Short, qPCR data collection guidance by Sam Palmer, and sample processing by the Vanderbilt VANTAGE Core.

