## Supplement for "Neuronal Modeling of Alternating Hemiplegia of Childhood Reveals Transcriptional Compensation and Replicates a Trigger-Induced Phenotype"

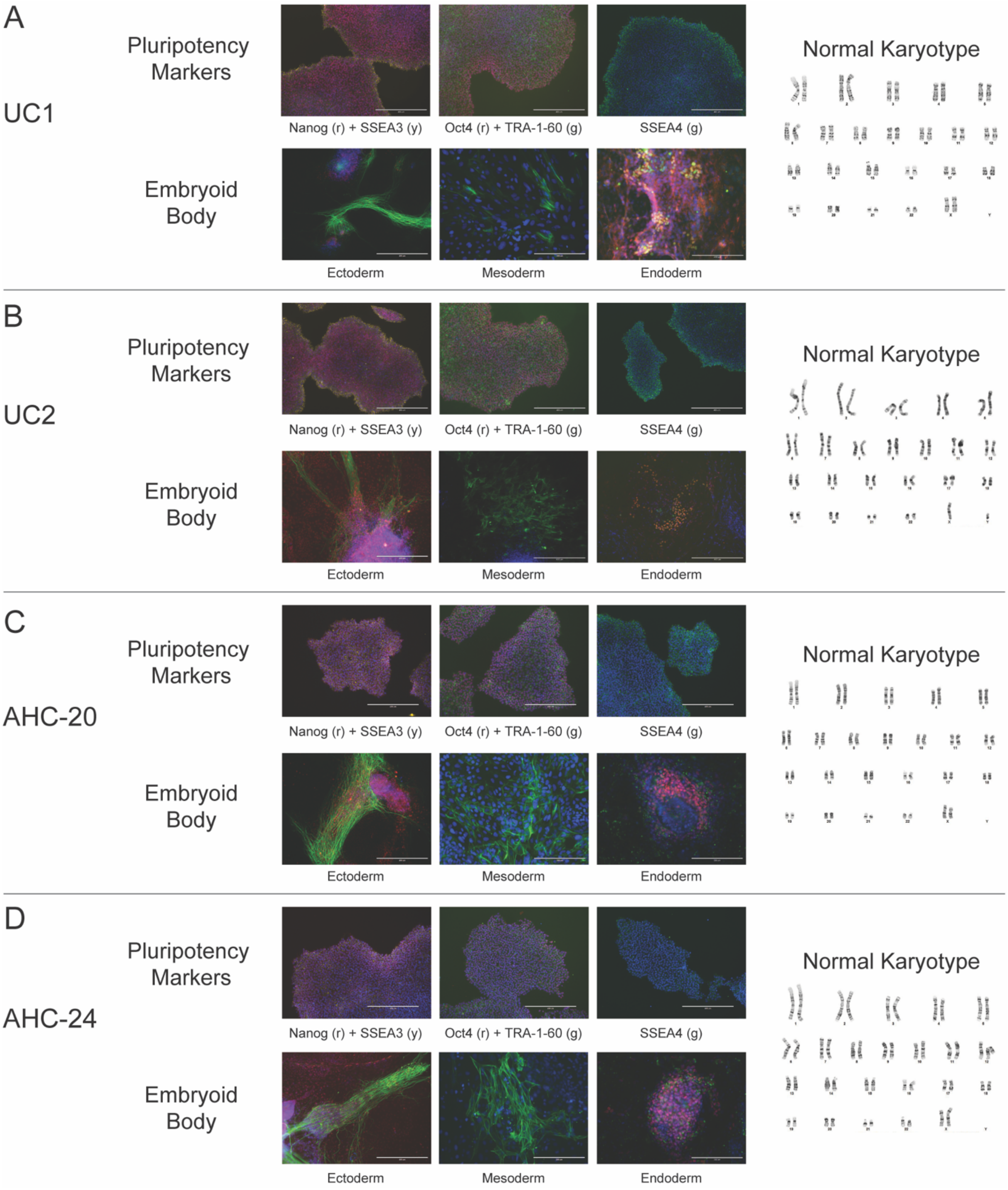
A
UC1
Pluripotency Markers
Nanog (r) + SSEA3 (y)
Oct4 (r) + TRA-1-60 (g)
SSEA4 (g)
Embryoid Body
Ectoderm
Mesoderm
Endoderm
Normal Karyotype
B
UC2
Pluripotency Markers
Nanog (r) + SSEA3 (y)
Oct4 (r) + TRA-1-60 (g)
SSEA4 (g)
Embryoid Body
Ectoderm
Mesoderm
Endoderm
Normal Karyotype
C
AHC-20
Pluripotency Markers
Nanog (r) + SSEA3 (y)
Oct4 (r) + TRA-1-60 (g)
SSEA4 (g)
Embryoid Body
Ectoderm
Mesoderm
Endoderm
Normal Karyotype
D
AHC-24
Pluripotency Markers
Nanog (r) + SSEA3 (y)
Oct4 (r) + TRA-1-60 (g)
SSEA4 (g)
Embryoid Body
Ectoderm
Mesoderm
Endoderm
Normal Karyotype

**Supplemental Figure 1: Validation of unrelated control and AHC patient iPSC lines.** Validation for the original clones used in this study (A) UC1, (B) UC2, (C) AHC-20, and (D) AHC-24. Pluripotency validation of iPSC clones for all lines included immunostaining for markers of pluripotency (Nanog, Oct4, SSEA4, SSEA3, and TRA-1-60). Differentiated stem cell aggregates (embryoid bodies) were also created and immunostained for characteristic markers of all three germline lineages: ectoderm (β-III-Tubulin [green] and Sox1 [red]), mesoderm (smooth muscle actin [green]), and endoderm (GATA4 [red], Sox17 [green]). Scale bars = 200 or 400 μm. DAPI nuclear stain shown in blue across all images. Metaphase spreads were performed (Genetic Associates, Nashville, TN) on all clones to ensure a normal karyotype. PCR analysis demonstrated that all lines had not incorporated the episomal pluripotency transduction plasmids into the host genome (not shown).

A

| | | |
|---|---|---|
| Clone IC1 | gRNA 1: TGCCATCTCACTGGCGTAC**A**(***AGG***)<br>ssODN Template-Induced Changes:<br>c.2443A>G - K815E correction to wildtype<br>c.2445G>A - E815E PAM-site synonymous | ssODN (1)<br>CTCCCCTCTCCGGCTCACCCGGCCTCCTCCGCCTAGGT<br>CCCTGCCATCTCACTGGCGTACG***AAG***CTGCCGAAAGCG<br>ACATCATGAAGAGACAGCCCAGGAACCCGCGGACGGAC<br>AAATTGGTC |
| Clone IC2 | gRNA 2: AGCCT**T**GTACGCCAGTGAGA(***TGG***)<br>Template-Induced Changes:<br>c.2443A>G - K815E correction to wildtype<br>c.2427C>A - A809A PAM-site synonymous | ssODN (2)<br>CCAGGAGGGTGGAGTCCTCCCCTCTCCGGCTCACCCGG<br>CCTCCTCCGCCTAGGTCCCTG***CAA***TCTCACTGGCGTACG<br>AGGCTGCCGAAAGCGACATCATGAAGAGACAGCCCAGG<br>AACCCGCGGACGGACAAATTGG |

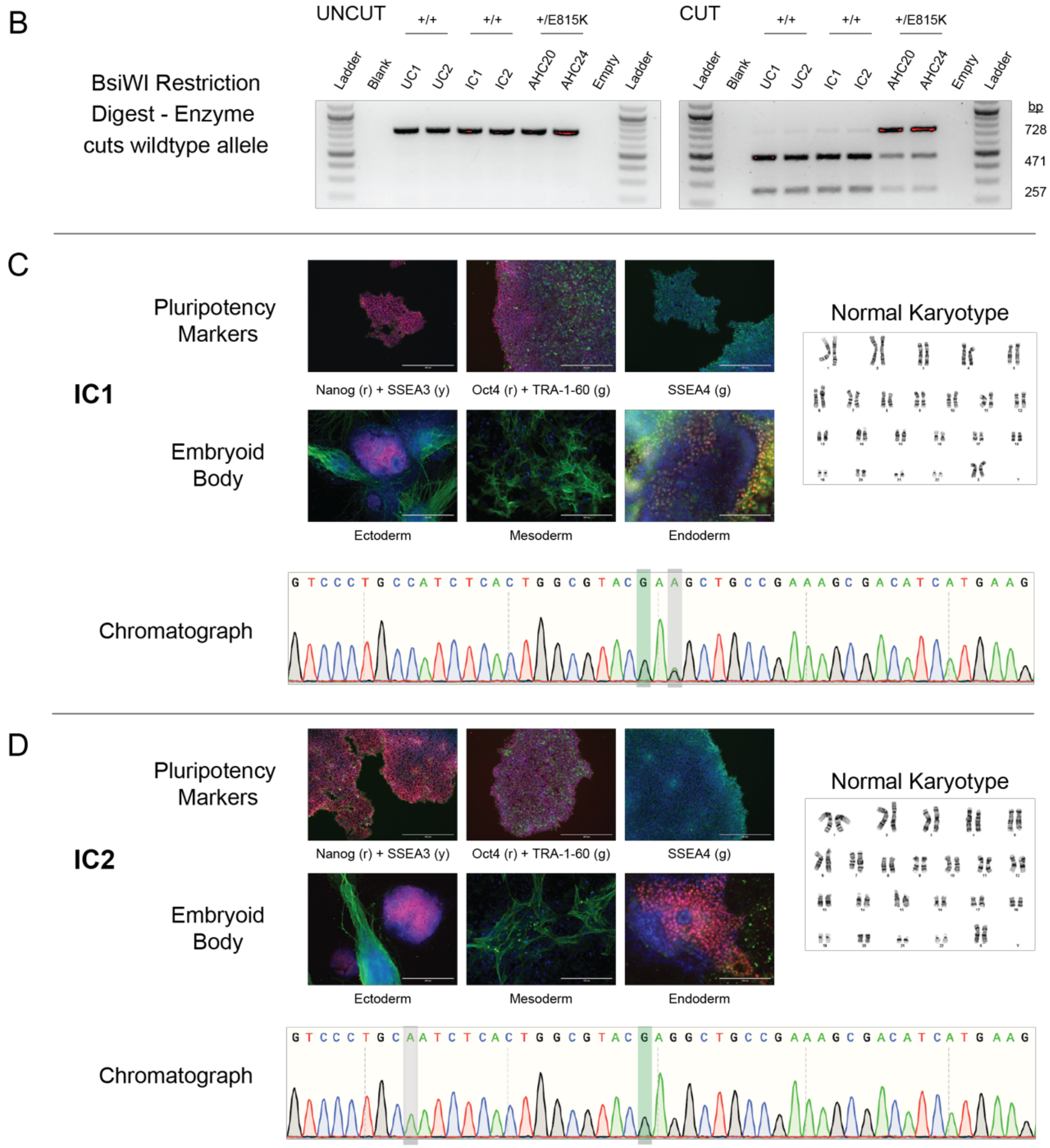

**Supplemental Figure 2: Isogenic correction strategy, confirmation, and pluripotency validation.** (A) Separate sgRNAs were designed to create two unique clones of isogenic corrected *ATP1A3*$^{+/+}$ lines from AHC patient line AHC-24. ssODN templates unique to each sgRNA included the intended c.2443A<G correction of the AHC mutation (green), along with synonymous changes to the PAM-site (red) to prevent repetitive cutting events by the Cas9 nuclease. (B) Following puromycin selection, surviving clones were screened using restriction enzyme BsiWI. The cut site for this enzyme is rescued with the corrected change. Multiple rounds of restriction enzyme screening occurred during cell culture to ensure the correct genotypes were used prior to neural differentiations. Shown here are date from the six iPSC clones used in the study. Sequencing of PCR products confirmed the sequence surrounding the cut site was unchanged outside of the intended alterations. Pluripotency validation was performed for isogenic corrected iPSCs as described in Figure S1 for (C) IC1 and (D) IC2 lines. For chromatographs, outlined green base represents correction to wildtype base location; synonymous PAM-site changes are highlighted in gray. For clone IC2, whole exome deep sequencing was performed to validate that sequences were homozygous corrected and not the result of uneven allelic amplification during PCR.

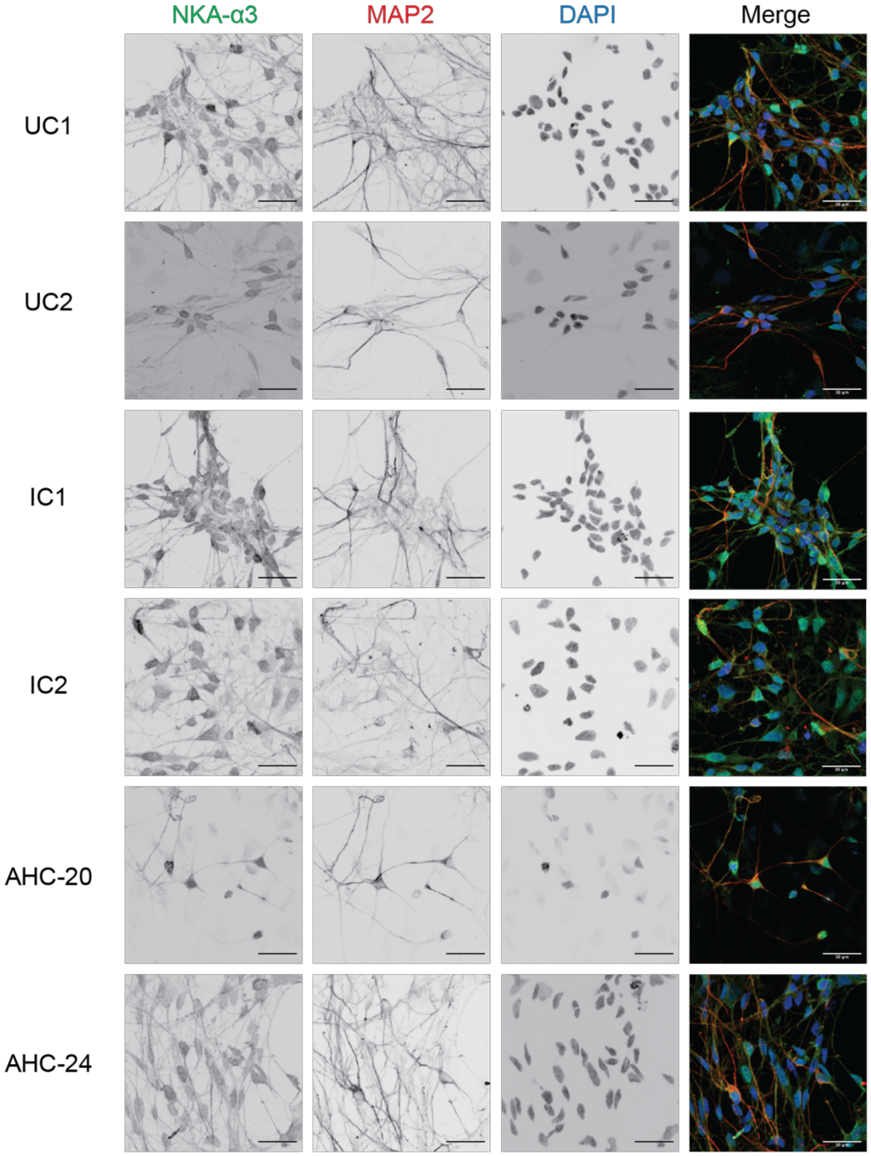


**Supplemental Figure 3: High resolution images of NKA-α3 expression in iPSC-derived neurons.** Confocal microscopy shows 60x magnification of NKA-α3 subunit (green) expression in MAP2 positive neurons (red) resulting from the Shi protocol at d60 of differentiation. DAPI shows nuclei (blue). No obvious differences in localization or expression patterns were noted across genotypes or differentiation by qualitative observation. Scale bar = 30 μm.

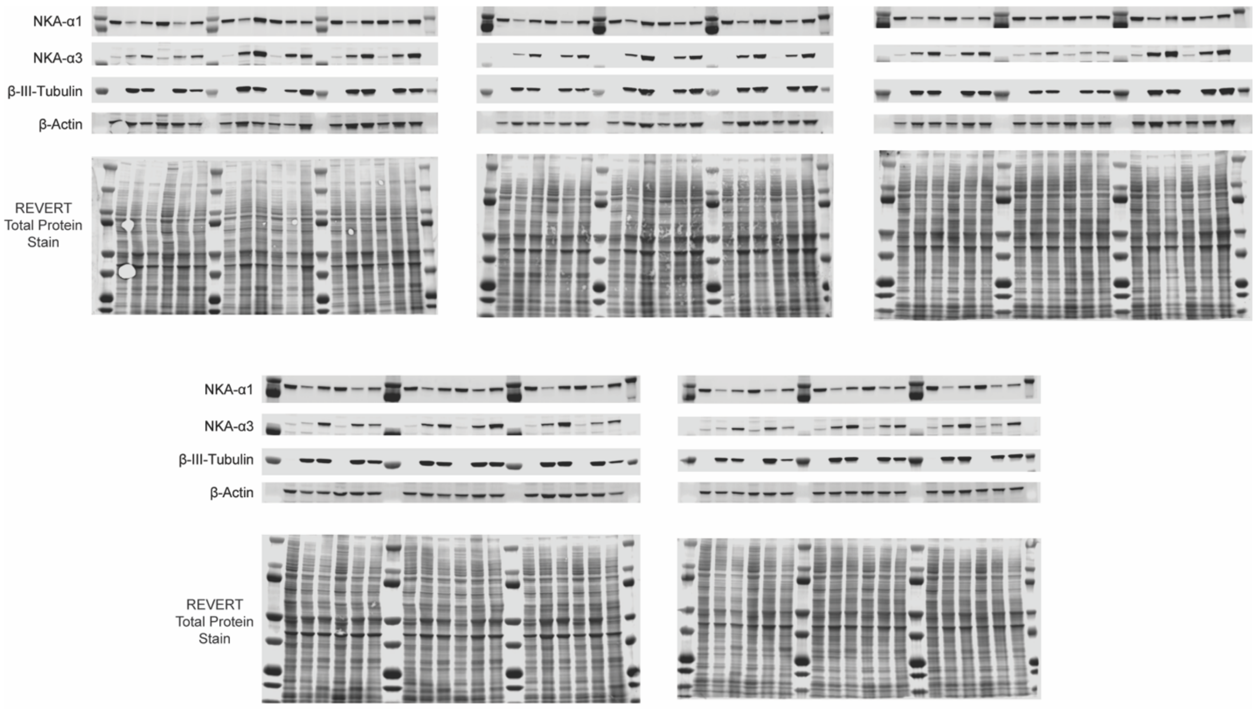


**Supplemental Figure 4: Full immunoblots and total protein stains used for normalization.** Blots were cut to size and probed for NKA-$\alpha$1, NKA-$\alpha$3, $\beta$-III-Tubulin, and $\beta$-Actin. $\beta$-Actin is shown as a visual guide for protein loading while intensity data was normalized to total protein shown here for each lane. Lysates from six clones of a given differentiation at three timepoints (d0, d32, d60) were run on the same gel, loaded as outlined in Figure 3.

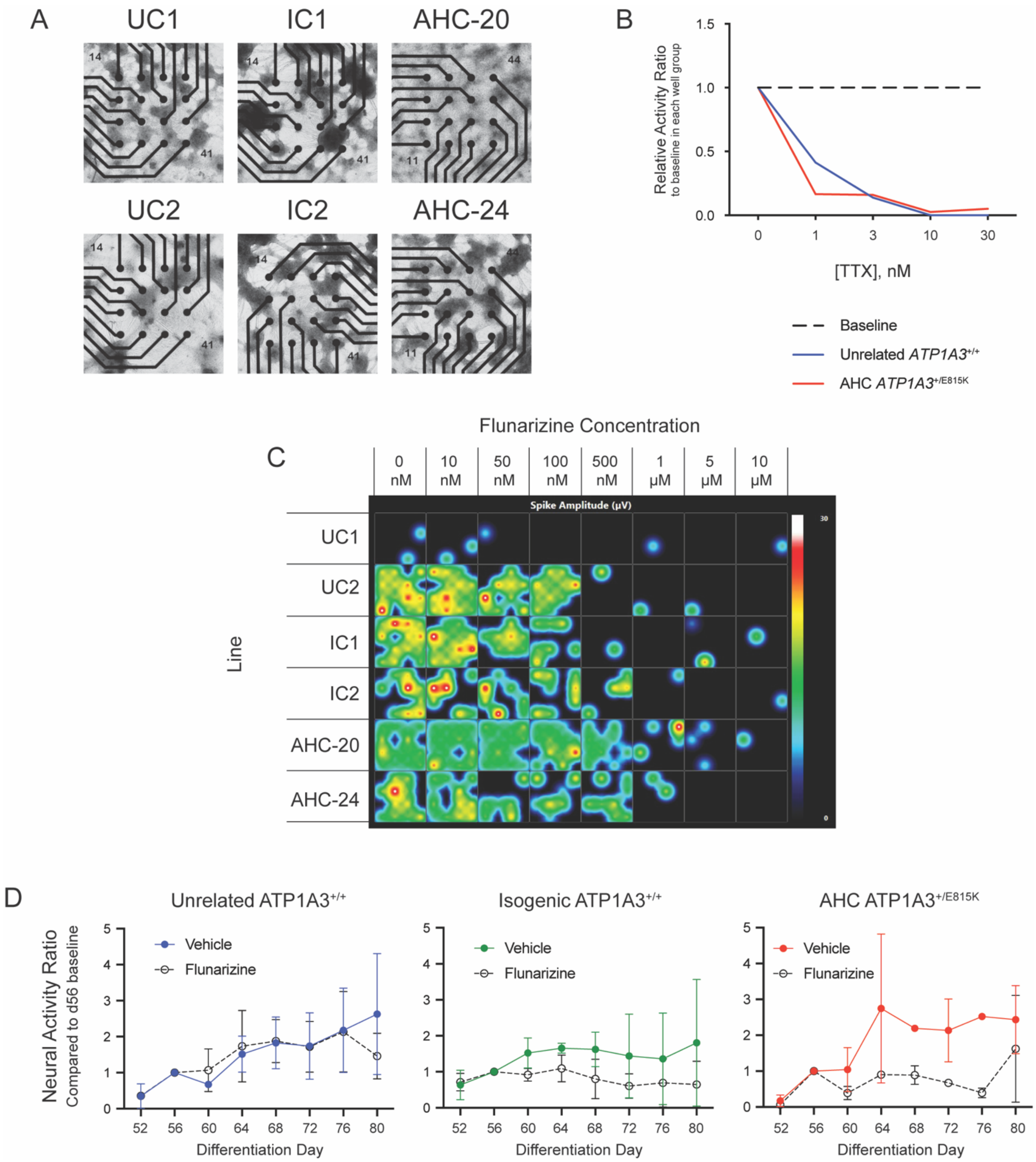


**Supplemental Figure 5: MEA plating: strategy, validation, and impact of flunarizine treatment on iPSC-derived neurons.** (A) Representative brightfield images of MEA plates at d80 shows clusters of neurons in relation to the 16 electrodes on each plate (black circles).

Following dispersion and plating at d45 of differentiation, neurons begin to cluster as seen in these photos. No obvious patterns of survival differences or overall network structures were noted between genotypes. (B) Treatment with increasing concentrations of tetrodotoxin (TTX) results in the elimination of activity in mixed cortical cultures, demonstrating the specificity of acquisition settings for neural spikes. (C) A heat map of spike amplitude shows inhibition of activity in the presence of high flunarizine concentrations. A chronic dosage concentration of 100 nM was chosen based on this dose-curve data and after literature review of applications in other model systems. (D) Flunarizine resulted in a loss of activity maturation over time in isogenic corrected and AHC cultures, but not in unrelated control neurons. Data represents mean activity for each clone summed across all baseline recording periods (n=2 for each data point), normalized to activity levels at d56 prior to treatment with vehicle or flunarizine. Error bars represent S.E.M.

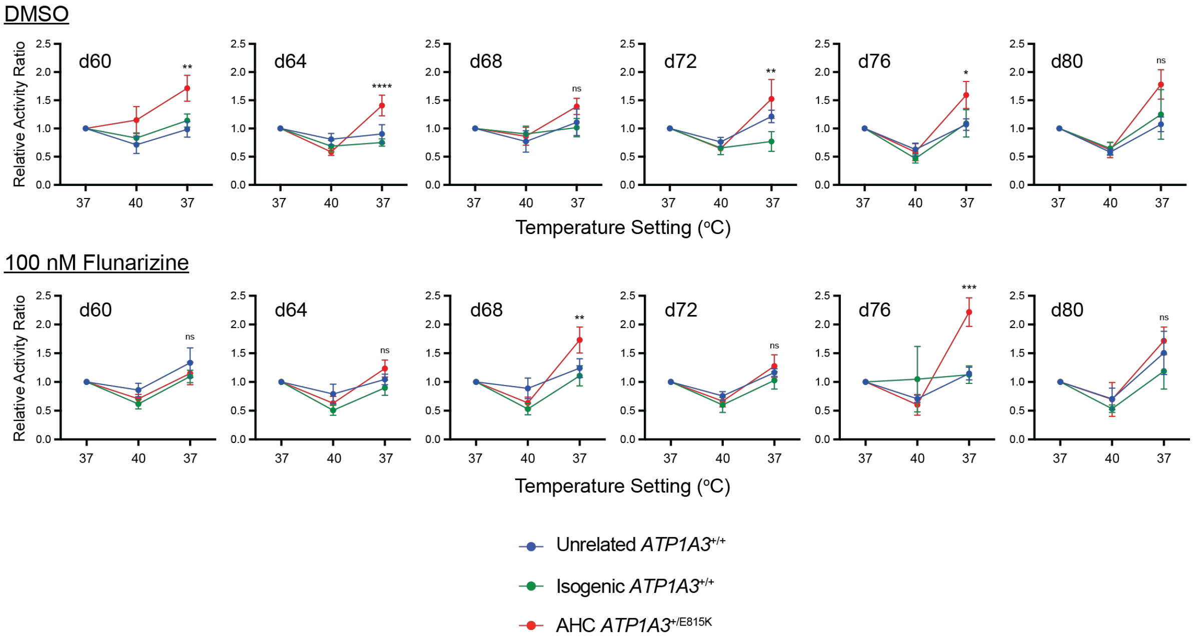


| | Sample Size (n) | | | | Significance (p-value) | | | | Significance (notation) | | | |
|---|---|---|---|---|---|---|---|---|---|---|---|---|
| | Day | UC-to-IC | AHC-to-UC | AHC-to-IC | Day | UC-to-IC | AHC-to-UC | AHC-to-IC | Day | UC-to-IC | AHC-to-UC | AHC-to-IC |
| **DMSO** | d60 | 7 | 9 | 7 | d60 | >0.9999 | 0.0013 | 0.0088 | d60 | ns | ** | ** |
| | d64 | 8 | 8 | 8 | d64 | 0.8382 | 0.0020 | <0.0001 | d64 | ns | ** | **** |
| | d68 | 8 | 9 | 10 | d68 | >0.9999 | 0.4820 | 0.1647 | d68 | ns | ns | ns |
| | d72 | 8 | 7 | 9 | d72 | 0.1708 | 0.4601 | 0.0037 | d72 | ns | ns | ** |
| | d76 | 9 | 6 | 7 | d76 | >0.9999 | 0.0135 | 0.0379 | d76 | ns | * | * |
| | d80 | 7 | 5 | 7 | d80 | >0.9999 | 0.0092 | 0.1112 | d80 | ns | ** | ns |
| | Sample Size (n) | | | | Significance (p-value) | | | | Significance (notation) | | | |
| | Day | UC-to-IC | AHC-to-UC | AHC-to-IC | Day | UC-to-IC | AHC-to-UC | AHC-to-IC | Day | UC-to-IC | AHC-to-UC | AHC-to-IC |
| **100 nM Flunarizine** | d60 | 7 | 7 | 6 | d60 | 0.5491 | 0.9586 | >0.9999 | d60 | ns | ns | ns |
| | d64 | 7 | 8 | 7 | d64 | 0.9455 | 0.6337 | 0.0720 | d64 | ns | ns | ns |
| | d68 | 8 | 7 | 8 | d68 | >0.9999 | 0.0352 | 0.0068 | d68 | ns | * | ** |
| | d72 | 8 | 7 | 8 | d72 | >0.9999 | >0.9999 | 0.3016 | d72 | ns | ns | ns |
| | d76 | 8 | 4 | 8 | d76 | >0.9999 | <0.0001 | 0.0007 | d76 | ns | **** | *** |
| | d80 | 8 | 4 | 5 | d80 | >0.9999 | >0.9999 | 0.5459 | d80 | ns | ns | ns |

**Supplemental Figure 6: Heat stress protocol MEA activity across recording days.** Neural spike activity ratios during heat stress protocol shown for each recording day, in the presence of vehicle or chronic flunarizine treatment. The included table describes the sample size along with significance values and notations for each data point. Data points represent values across 5 differentiations with 2 clones per genotype. Significance notations represent AHC-to-isogenic comparisons in the recovery period (37C). As this data is more granular and sample sizes are variable due to exclusion criteria, significance for 2-way ANOVA with Bonferroni's multiple comparisons test is shown as significant at $p < 0.05$ (*), $p > 0.01$ (**), $p<0.001$ (***), and $p<0.0001$ (****); ns $p > 0.05$. Error bars represent S.E.M.

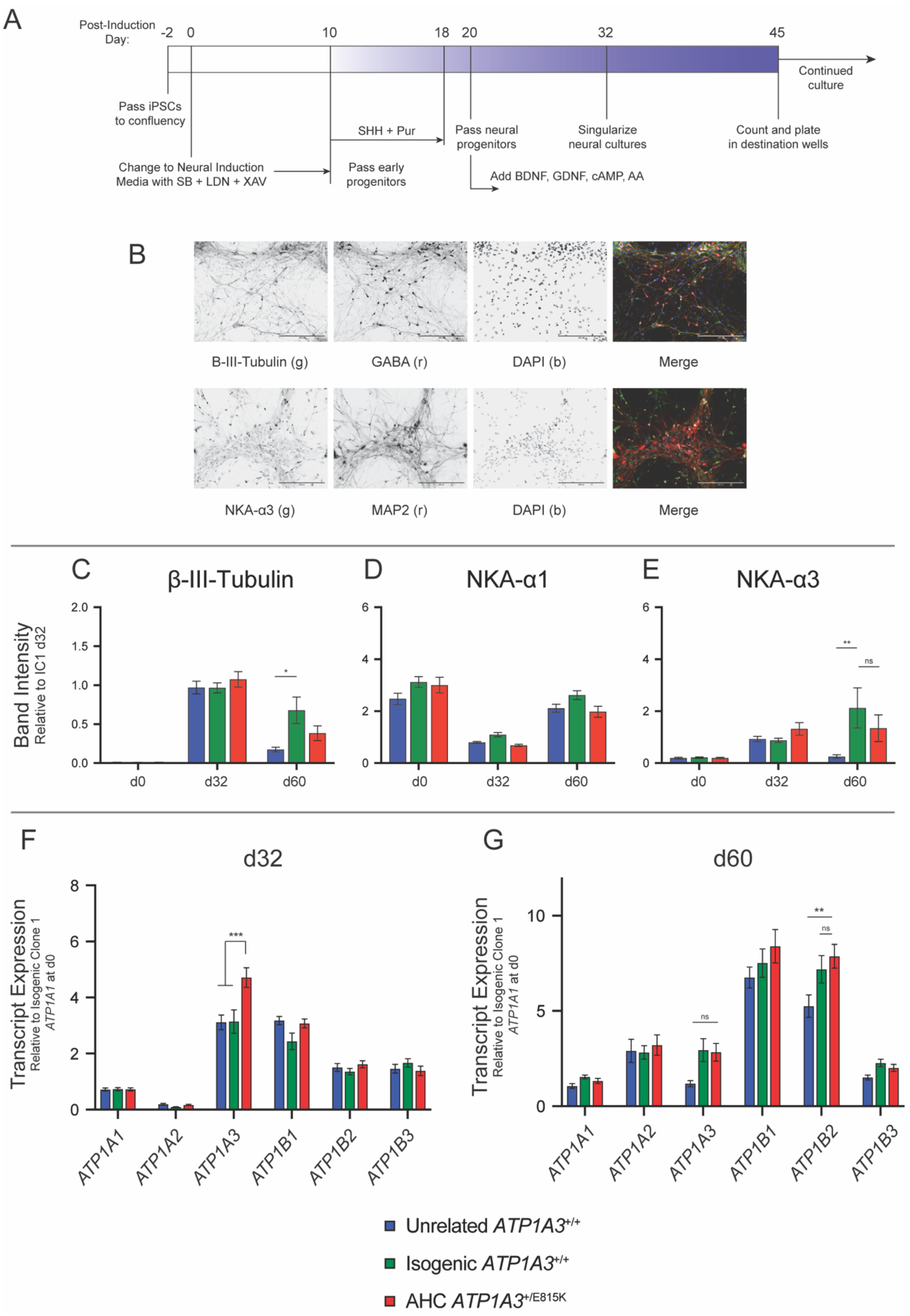
A
Post-Induction Day:
-2 0 10 18 20 32 45
Continued culture
Pass iPSCs to confluency
Change to Neural Induction Media with SB + LDN + XAV
SHH + Pur
Pass early progenitors
Pass neural progenitors
Add BDNF, GDNF, cAMP, AA
Singularize neural cultures
Count and plate in destination wells
B
B-III-Tubulin (g)
GABA (r)
DAPI (b)
Merge
NKA-α3 (g)
MAP2 (r)
DAPI (b)
Merge
C
β-III-Tubulin
D
NKA-α1
E
NKA-α3
Band Intensity
Relative to IC1 d32
d0
d32
d60
F
d32
G
d60
Transcript Expression
Relative to Isogenic Clone 1
ATP1A1 at d0
ATP1A1
ATP1A2
ATP1A3
ATP1B1
ATP1B2
ATP1B3
Unrelated ATP1A3+/+
Isogenic ATP1A3+/+
AHC ATP1A3+/E815K

**Supplemental Figure 7: GABAergic predominant neuronal differentiation and NKA subunit expression patterns generally replicate results from mixed cortical differentiation.** (A) A different protocol (Maroof et al., 2013) was utilized to test the specificity of our results to a single neural differentiation scheme. Utilizing Wnt-antagonism and SHH agonism in addition to standard dual-SMAD inhibition techniques, this protocol produces significantly higher proportions of GABAergic neurons in culture as described in the original protocol. (B) Neurons derived from this protocol express standard neural markers including β-III-Tubulin and MAP2 when fixed at d60. GABAergic staining shows increased amounts of GABA-positive cells that also stain positive for the NKA-α3 subunit. DAPI shown as a nuclear stain; scale bars = 200 μm. (C) Immunoblotting data shows an increase in β-III-Tubulin expression at d32 that is nonsignificant between genotypes. Due to the presence of contaminating cell types in this protocol by d60 of differentiation, β-III-Tubulin and other neural markers become less consistently expressed resulting in spurious differences (d60 isogenic-to-unrelated $p = 0.0001$). (D) Expression of the NKA-α1 subunit follows a very similar pattern to mixed cortical differentiation with nonsignificant differences at individual timepoints between genotypes. (E) No significant differences in NKA-α3 subunit protein expression were noted between isogenic corrected and AHC lines across GABAergic shifted differentiation. (F) qPCR analysis showed a significant elevation of *ATP1A3* transcripts at d32 ($p < 0.0001$) in AHC iPSC-derived neurons compared to both control groups. (G) By d60, deterioration of neural purity resulted in a relative loss of *ATP1A3* transcript levels compared to d32 with nonsignificant differences between isogenic control and AHC cultures. n = 10 (5 differentiations with 2 clones per genotype). 2-way ANOVA with Bonferroni's multiple comparisons test; significant findings notated by: * $p < 0.005$, ** $p < 0.001$, *** $p < 0.0001$; ns = not significant ($p > 0.01$).

**Primers for Genotyping & Validation (Integrated DNA Technologies)**

| Primer Name | Primer Sequence |
|---|---|
| *ATP1A3* Exon 18 PCR Forward Primer | CGCCTGATCTTCGACAACCT |
| *ATP1A3* Exon 18 PCR Reverse Primer | TAACCTGGAGCCCCTCTCTC |
| *ATP1A3* Exon 18 PCR Nested Forward Primer | TACACCCTGACCAGCAATATC |
| PX459 U6P Primer for Ligation Validation | GGACTATCATATGCTTACCG |

**Primers and Probes for qPCR (Applied Biosystems)**

| Transcript | Assay Identifier |
|---|---|
| *GAPDH* | Hs99999905_m1 |
| *ATP1A1* | Hs00167556_m1 |
| *ATP1A2* | Hs00265131_m1 |
| *ATP1A3* | Hs00958036_m1 |
| *ATP1B1* | Hs00426868_g1 |
| *ATP1B2* | Hs01020302_g1 |
| *ATP1B3* | Hs00740857_mH |

**Antibodies for Immunostaining and Immunoblotting**

| Antibody Target | Application | Host | Dilution | Provider | Catalog # |
|---|---|---|---|---|---|
| Nanog | ICC | Rabbit | 1:250 | Cell Signaling | 4903 |
| Oct4 | ICC | Rabbit | 1:250 | Cell Signaling | 2750 |
| TRA-1-60 | ICC | Mouse | 1:250 | EMD Millipore | MAB4360 |
| SSEA3 | ICC | Rat | 1:250 | EMD Millipore | MAB4303 |
| SSEA4 | ICC | Mouse | 1:250 | EMD Millipore | MAB4304 |
| β-III-Tubulin | ICC / WB | Mouse | 1:250 / 1:2000 | EMD Millipore | MAB1637 |
| Sox1 | ICC | Rabbit | 1:100 | Genetex | gtx62974 |
| Smooth Muscle Actin | ICC | Mouse | 1:200 | Abcam | ab5694 |
| GATA4 | ICC | Rabbit | 1:200 | Abcam | ab84593 |
| Sox17 | ICC | Mouse | 1:100 | Abcam | ab84990 |
| GABA | ICC | Rabbit | 1:500 | Sigma | a2052 |
| NKA-α3 | ICC | Mouse | 1:200 | Thermo Fisher | MA3-915 |
| NKA-α3 | WB | Rabbit | 1:1000 | Abcam | ab182572 |
| NKA-α1 | WB | Mouse | 1:250 | Iowa DSHB | A6F |
| β-Actin | WB | Rabbit | 1:2000 | Cell Signaling | 4967S |

**Reagent Lists**

iPSC Culture & Maintenance

| Reagent / Product | Provider | Catalog # |
|---|---|---|
| mTeSR1 Complete Kit | Stemcell Technologies | 85875 |
| Matrigel Matrix | Corning | 356234 |
| ReLeSR | Stemcell Technologies | 05872 |

Neural Differentiation & Maintenance Reagents

| Reagent / Product | Provider | Catalog # |
|---|---|---|
| Accutase | Innovative Cell Technologies | AT-104 |
| Ascorbic Acid | Sigma | A4403 |
| B27 Supplement | Gibco | 17504044 |
| B27 Plus Supplement | Gibco | A3582801 |
| BDNF | Peprotech | 450-02 |
| BME | Sigma | M6250 |
| Dibutyryl cAMP | Sigma | D0627 |
| Dispase | Stemcell Technologies | 07923 |
| DMEM | Gibco | 11995-065 |
| DMEM/F12 + Glutamax | Gibco | 10565-018 |
| FGF2 | Peprotech | 100-18B |
| GDNF | Peprotech | 450-10 |
| Glutamax | Gibco | 35050061 |
| IGF-1 | Peprotech | 100-11 |
| Insulin | Life Technologies | 12585014 |
| KnockOut Serum Replacement | Gibco | 10828028 |
| LDN-189193 | Tocris | 6053 |
| N2 Supplement | Gibco | 17502048 |
| Neurobasal Medium | Gibco | 21103049 |
| Non-essential amino acids | Sigma | M7145 |
| Penicillin-Streptomycin | Gibco | 15140122 |
| Purmorphamine | EMD Millipore | 540220 |
| rhSHH (C24II) | R&D Systems | 1845-SH |
| SB-431542 | Cayman Chemical | 13031 |
| XAV939 | Cayman Chemical | 13596 |
| Y-27632 (ROCK-i) | Cayman Chemical | 10005583 |

**Neural Differentiation: Base Media Recipes**

Neural Maintenance Media (NMM, 500 mL)

- 242 mL Neurobasal
- 242 mL DMEM/F12 + Glutamax
- 2.5 mL Non-Essential Amino Acids
- 3.75 mL 100x Glutamax
- 5.0 mL Penicillin-Streptomycin
- 2.5 mL 100x N2 Supplement
- 5.0 mL 50x B27 Supplement
- 312.5 μL Insulin (2.5 μg/mL)
- 1.75 μL BME (50 μM)

Knockout Serum Replacement Media (KSR, 300 mL)

- 252 mL DMEM
- 45 mL KSR
- 3 mL Penicillin-Streptomycin
- 1.05 μL BME (50 μM)

Mixed Cortical Neural Differentiation Media (mcNDM, 500 mL)

- 475 mL Neurobasal
- 10 mL 50x B27 Supplement
- 5 mL 100x N2 Supplement
- 5 mL Non-Essential Amino Acids
- 5 mL Pen-Strep
- Supplemented day of feeding with:
    - 10 ng/mL BDNF
    - 10 ng/mL GDNF
    - 10 ng/mL IGF-1
    - 1 μM Dibutyryl cAMP

GABAergic Neural Differentiation Media (gNDM, 500 mL)

- 475 mL Neurobasal
- 10 mL 50x B27 Supplement
- 5 mL 100x Glutamax
- 5 mL Non-Essential Amino Acids
- 5 mL Pen-Strep
- Supplemented day of feeding with:
    - 10 ng/mL BDNF
    - 10 ng/mL GDNF
    - 200 μM Ascorbic Acid
    - 200 μM Dibutyryl cAMP
